# Controllable Protein Design via Autoregressive Direct Coupling Analysis Conditioned on Principal Components

**DOI:** 10.1101/2025.08.18.669886

**Authors:** Francesco Caredda, Lisa Gennai, Paolo De Los Rios, Andrea Pagnani

## Abstract

We present FeatureDCA, a statistical framework for protein sequence modeling and generation that extends Direct Coupling Analysis (DCA) with biologically meaningful conditioning. The method can leverage different kinds of information, such as phylogeny, optimal growth temperature, enzymatic activity or, as in the case presented here, principal components derived from multiple sequence alignments, and use it to improve the learning process and consequently efficiently condition the generative process. FeatureDCA allows sampling to be guided toward specific regions of sequence space while maintaining the efficiency and interpretability of Potts-based inference. Across multiple protein families, our autoregressive implementation of FeatureDCA matches or surpasses the generative accuracy of established models in reproducing higher-order sequence statistics while preserving substantial sequence diversity. Structural validation with AlphaFold and ESMFold confirms that generated sequences adopt folds consistent with their intended wild-type targets. In a detailed case study of the Response Regulator family (PF00072), which comprises distinct structural subclasses linked to different DNA-binding domains, FeatureDCA accurately reproduces class-specific architectures when conditioned on subtype-specific principal components, highlighting its potential for fine-grained structural control. Predictions of experimental deep mutational scanning data show accuracy comparable to that of unconditioned autoregressive Potts models, indicating that FeatureDCA also captures local functional constraints. These results position FeatureDCA as a flexible and transparent approach for targeted sequence generation, bridging statistical fidelity, structural realism, and interpretability in protein design.

## Introduction

The ability to generate novel functional protein sequences is a central challenge in computational biology and protein design. During the past decade, statistical models trained on evolutionary data, particularly those derived from multiple sequence alignments (MSAs), have demonstrated remarkable success in capturing the statistical, structural, and functional constraints that shape natural protein families. Among these, models based on the Potts framework, such as Boltzmann machine Direct Coupling Analysis (bmDCA)[1, 2, 3, 4] and autoregressive DCA (ArDCA)[5], have established themselves as powerful tools for inferring residue-residue interactions, predicting mutational effects, and designing synthetic sequences with natural-like properties [6].

Despite their success, a fundamental limitation of these models lies in their lack of control over the output distribution. Once trained, sampling from such models yields sequences that broadly match the statistical patterns of the training MSA, but offers little to no ability to steer generation toward user-defined features or functional subtypes. In practice, however, protein design often demands precisely this kind of control, whether to target a specific structural class, functional behavior, or region in sequence space.

Building on advances in sequence modeling, several generative approaches have emerged in recent years for protein sequence design, including variational autoencoders (VAEs) [7, 8], generative adversarial networks (GANs)[9], protein language models (PLMs) [10, 11, 12, 13, 14, 15, 16, 17], and diffusion-based methods [18]. Among these, DeepSequence [7] was an early VAE model that demonstrated strong performance in mutational effect prediction and unsupervised modeling of protein families. More recent protein language models, such as ESM [11, 19, 20] and ProGen [10], leverage large-scale transformer architectures trained on millions of unaligned sequences to learn general-purpose protein representations. These models have shown impressive capabilities in sequence generation, mutational scanning, and even zero-shot structure prediction. While flexible and expressive, such models often require massive datasets and computational resources, and they generally lack interpretability and direct access to the coevolutionary statistics encoded in MSAs. Moreover, their mechanisms for guiding sequence generation, such as prompt tuning or conditioning through templates, are often indirect or non-transparent, and do not solve the fundamental problem of explicitly conditioning generation on user-defined features within a statistical framework. In contrast, DCA-based models operate directly on aligned homologous sequences, offering a principled, data-efficient, and interpretable alternative that tightly reflects evolutionary constraints and which, with the right extensions, can be made controllable.

In this work, we introduce FeatureDCA, a principled extension of the DCA framework that enables conditional generation of protein sequences. Our approach incorporates low-dimensional, biologically meaningful features, particularly the top principal components of the MSA, as conditioning inputs to the model. The inference and sampling are performed in an autoregressive manner, embedding the co-evolutionary data and the low-dimensional features into autoregressive conditional distributions to guide generation toward specific points or regions in feature space, enabling a new level of controllability and interpretability in statistical protein design. A conceptually similar approach underlies AF-Cluster [21], which shows that clustering sequences within a single MSA can separate structural signals associated with different conformations. While both approaches exploit this separation, FeatureDCA incorporates it directly into a generative model, enabling continuous conditioning and targeted sequence design.

We demonstrate that FeatureDCA achieves generative accuracy comparable to or better than state-of-the-art models like bmDCA and ArDCA, while also enabling controllable, feature-conditioned sampling. Using PCA-based conditioning, FeatureDCA generates synthetic sequences that accurately reproduce the statistical properties of natural MSAs, such as pairwise correlations and principal component distributions, while maintaining substantial sequence diversity, as measured by Hamming distance. This balance between statistical fidelity and diversity is crucial for effective protein design, where novel sequences must respect evolutionary constraints but also explore unseen regions of sequence space.

In addition to statistical validation and sequence-level metrics, we assess the structural realism of generated sequences using AlphaFold [22, 23] and ESMFold [19], showing that FeatureDCA maintains high structural consistency with experimentally determined wild-type folds. As a biologically grounded case study, we focus on the Response Regulator (RR) family (Pfam ID PF00072), a large and diverse set of bacterial proteins involved in signal transduction. RR proteins share a conserved receiver domain but exhibit distinct dimerization architectures depending on their DNA-binding domains. We show that FeatureDCA, when conditioned on PCA-derived features, can selectively generate sequences corresponding to different RR subclasses, illustrating its capacity to navigate biologically meaningful variation and control functional outcomes in sequence design. Finally, we also evaluate the model’s predictive power on experimental deep mutational scanning (DMS) data, demonstrating competitive performance in capturing the functional effects of single-residue mutations.

A schematic overview of the FeatureDCA framework, including training on aligned MSAs, conditioning on PCA-derived features, and downstream applications such as conditional sampling and fitness prediction, is provided in Figure 1. Together, our results position FeatureDCA as a powerful and flexible framework for generative modeling in protein sequence space, combining the strength of Potts-based inference with the adaptability of feature-based conditioning.

**Figure 1:**
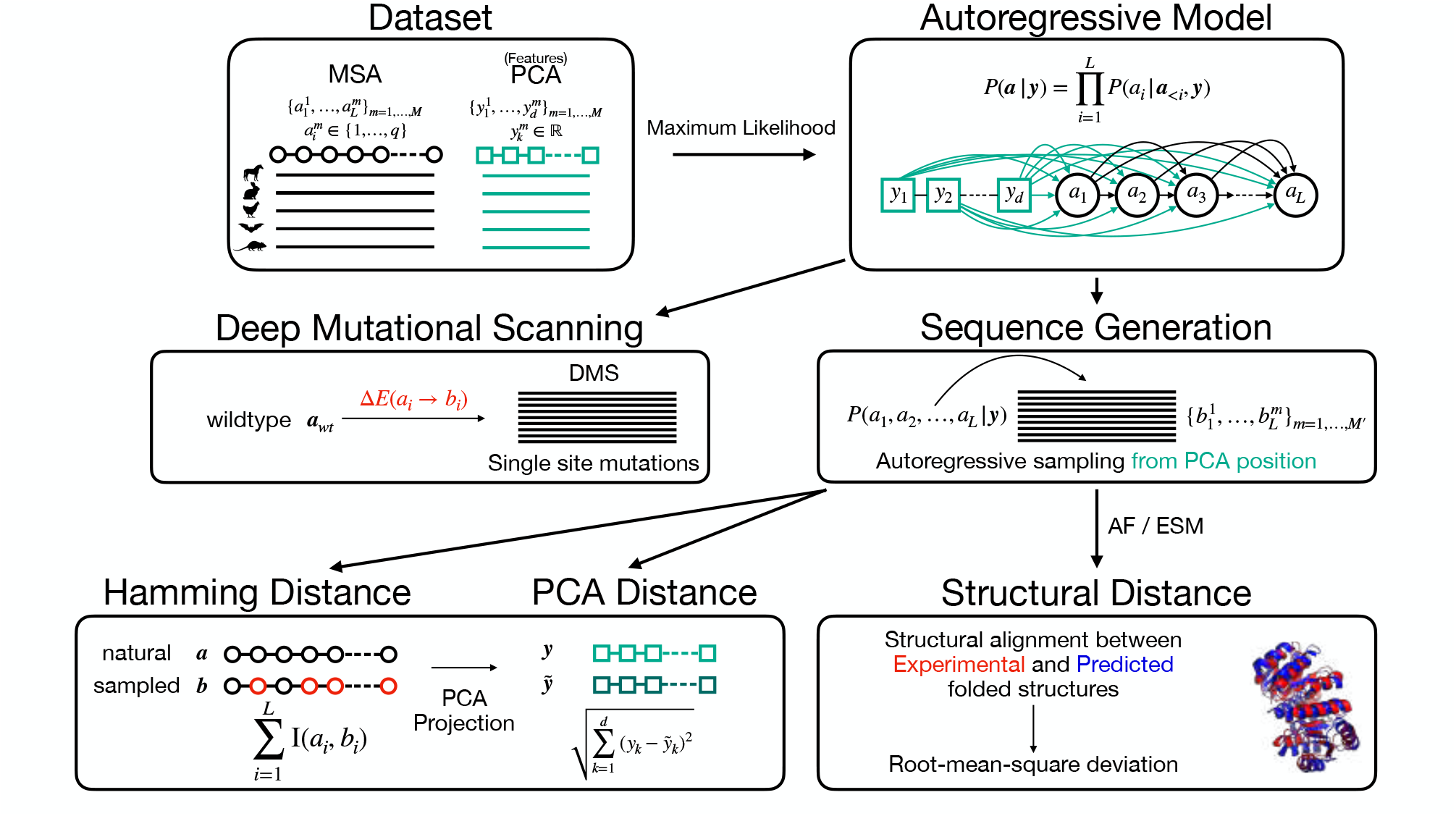
Overview of the generative modeling pipeline. A multiple sequence alignment (MSA) of homologous sequences is embedded using PCA to produce continuous feature vectors for each sequence. An autoregressive model is trained to learn the conditional distribution *P* (*a*_1_, …, *a*_*L*_ |***y***), where ***y*** encodes the PCA features. The model is used for multiple downstream tasks: (1) Deep Mutational Scanning (DMS) prediction via single-site substitutions and computation of energy changes Δ*E*(*a*_*i*_ → *b*_*i*_); (2) Sequence generation by autoregressive sampling conditioned on ***y***; and (3) Evaluation of generated sequences by comparing them to natural sequences using Hamming distance, PCA embedding distance, and structural RMSD between predicted and experimental structures.

## Results

### Architecture

We present a novel extension of the standard autoregressive Direct Coupling Analysis (ArDCA) method [5], in which sequence-dependent vectors of biologically relevant features are embedded in the amino acid space of a protein family, thereby constraining the sampling of new sequences to specific, user-defined characteristics. Starting from the exact decomposition

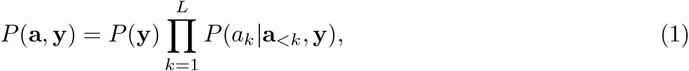

we model the single-site distribution of amino acid *a*_*k*_, conditioned on amino acids **a**_*<k*_ = {*a*_1_, *a*_2_, …, *a*_*k−*1_} and feature vector ***y*** ∈ ℝ^*d*^ as a Boltzmann measure over the energy function

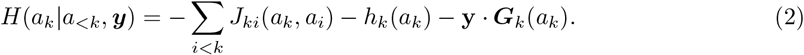

Here, the first two terms represent the standard Potts model that captures co-evolution and conservation signals through an interaction tensor *J*_*ki*_(*a*_*k*_, *a*_*i*_) and a local field *h*_*k*_(*a*_*k*_) [24, 25]. The final term couples the vector of external conditioning characteristics ***y*** with an embedding ***G***_***k***_(*a*_*k*_) of the amino acid in the feature space. This formulation effectively combines evolutionary constraints with additional design conditions, offering a robust, flexible framework for generative protein design. A detailed derivation of this functional form, based on a Gaussian approximation of the conditional distribution, is provided in Supporting Information (SI) Section 2.

The inference of the model’s parameters is performed by an L2-regularized maximum likelihood estimation over a joint dataset of a multiple sequence alignment (MSA) and its corresponding conditioning features:

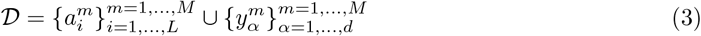

with 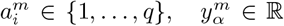, where *L* is the length of the multiple sequence alignment, *M* the number of MSA sequences, *d* is the dimension of the feature vector, and *q* is the alphabet size of MSA (*q* = 21 for proteins: 20 amino acids plus the gap symbol, see SI Sections 1 and 3 for implementation details). Given the model parameters, the sampling is performed in an autoregressive fashion. Starting from a given feature vector, amino acids are sampled iteratively, conditioned on the previous positions along the chain. This is a highly efficient sampling method, as it consists of *L* different samples of univariate distributions of the form:

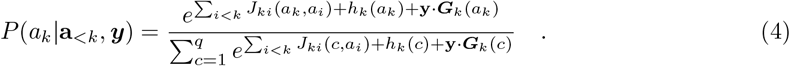

Motivated by the need to address sequence generation in specific regions of the Principal Component (PC) space, along with the simplicity of performing a Principal Component Analysis (PCA) of an MSA, we apply our model to the case in which the feature vectors represent the first *d* PCs of each sequence (here and in the following we assume that PCAs dimensions are ordered in decreasing variance of the data). However, the nature of the feature vector can be heterogeneous: it can represent different biological traits, such as structural or functional characteristics, as well as evolutionary or experimental ones. Examples of these can be optimal growth temperatures, folding temperatures, binding specificity, phylogenetic profiles, stability, or mutational fitness.

### Generativity

The minimal requisite of the model is to sample sequences that are statistically indistinguishable from the natural ones. This means reproducing both the pairwise frequency statistics and the PCA projection of the natural MSA. As a metric for the pairwise frequency statistics, we use the Pearson coefficient of the connected correlations of the natural and generated MSAs:

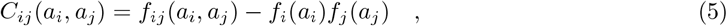

where *f*_*i*_ and *f*_*ij*_ are the single- and pair-wise frequency counts of the alignment, see SI Section 1 for details. To quantitatively compare the similarities between the PC projections of the natural and generated datasets, we introduce the entropic-regularized Wasserstein distance, also known as Sinkhorn divergence or Earth Mover’s distance [26]. This metric defines the optimal transport distance between two distributions. Technical details and definitions can be found in SI Section 5. As a general benchmark for the generative capabilities of the model, we use bmDCA and standard ArDCA, which are considered the state-of-the-art architectures in terms of accuracy and computational efficiency, respectively. Relative to protein family PF13354 (beta-lactamase), Figure 2A) represents the PCA distribution of the natural MSA compared to those of the MSAs sampled by bmDCA, ArDCA, and FeatureDCA. In particular, each sequence from FeatureDCA is generated conditioning on a unique *d*-dimensional point in the PCA projection of the natural distribution. In Figure 2A), FeatureDCA is trained with *d* = 2 principal components, which is enough to reproduce a PCA distribution qualitatively and quantitatively indistinguishable from that of bmDCA.

**Figure 2:**
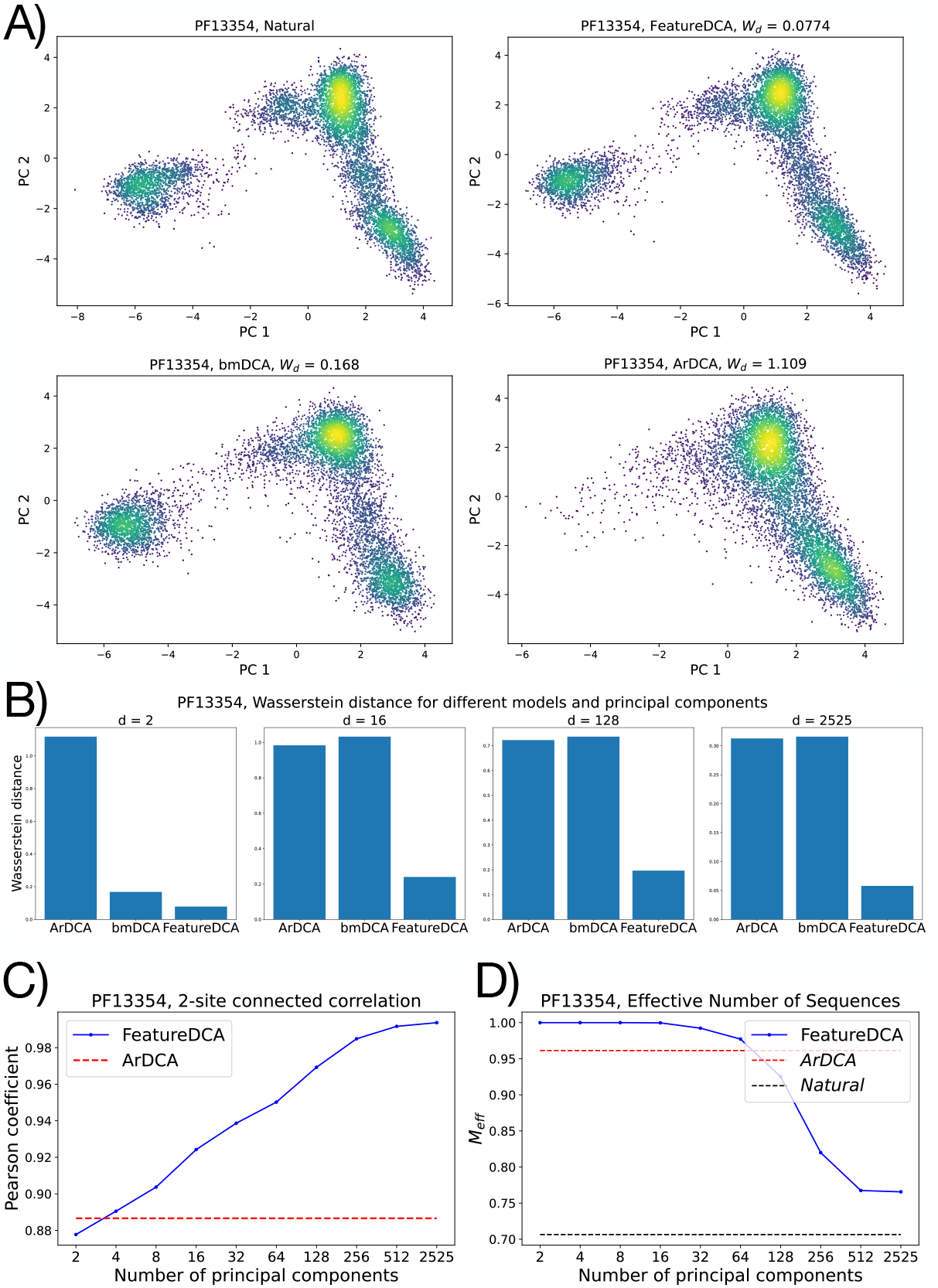
**A)** Projection onto the first two principal components of natural and generated MSAs from protein family PF13354. **B)** Wasserstein distance between the *d*-dimensional PCA distributions of the natural and generated MSA for different models. FeatureDCA was trained with the corresponding number of principal components. **C)** Pearson coefficient of the 2-site connected correlation between natural and generated data as a function of the number of principal components learned during training. **D)** Effective depth percentage of the generated dataset as a function of the number of principal components learned during training.

Figure 2B) shows the Wasserstein distance between the natural and generated datasets when projected in the *d*-dimensional PCA space for different generative models. In particular, the PCA distribution of data generated from FeatureDCA can accurately reproduce the first *d* components, provided they have been learned during training, while bmDCA and ArDCA do not reproduce components higher than the second, see SI Section 6. Figure 2C) represents the Pearson coefficient of the connected correlation defined in Equation 5 between natural and generated data. As the number of principal components increases, so does the Pearson coefficient relative to FeatureDCA, outperforming either ArDCA or bmDCA. However, when the number of principal components approaches *d*_max_ = 2525, representing the number of components required to explain 99% of the variance of the MSA, the model overfits the data and generates sequences that are identical to the natural ones. In this regime, the model begins to memorize natural sequences rather than generalizing, as indicated by near-zero Hamming distances to training data. Finally, Figure 2D shows the behavior of the effective depth *M*_eff_, the number of effectively unique sequences as defined in SI Section 1, as a function of the number of principal components used during training.

#### In-place generativity

Although the model can globally reproduce the PC projection to the point of improving the generativity of the most reliable models, such as bmDCA, the actual goal of the architecture is to sample sequences at specific points in the PC space. Given a wild-type sequence ***a***^*wt*^ located at point ***y***^*wt*^ ∈ ℝ^*d*^ in the PC space, sequences generated conditioned on that point should be comparable to the wild type in terms of Hamming distance and Euclidean distance in the PC space. In Figure 3, we compare the average Hamming and Euclidean distances of samples of a thousand sequences generated conditionally to three different wild-type sequences as a function of the number of principal components learned during training and used for the sampling. It can be seen that, using fewer principal components, the generated sequences are spread on average over a cloud centered around the corresponding wild type, with a Hamming distance between 60% and 80%. As the number of principal components increases, the average position of each sample cluster shifts toward its target wild type, while the variance of the distribution decreases. Accordingly, the average Hamming distance between samples and non-corresponding wild types approximates the actual Hamming dis-tance between different wild types. However, when the number of principal components approaches *d*_max_, the Hamming distance between a sample and its corresponding wild type goes to zero, highlighting the memorization of the natural dataset due to overfitting. From a statistical point of view, a number of principal components between 32 and 128 results in a good trade-off between generative accuracy and generalization.

**Figure 3:**
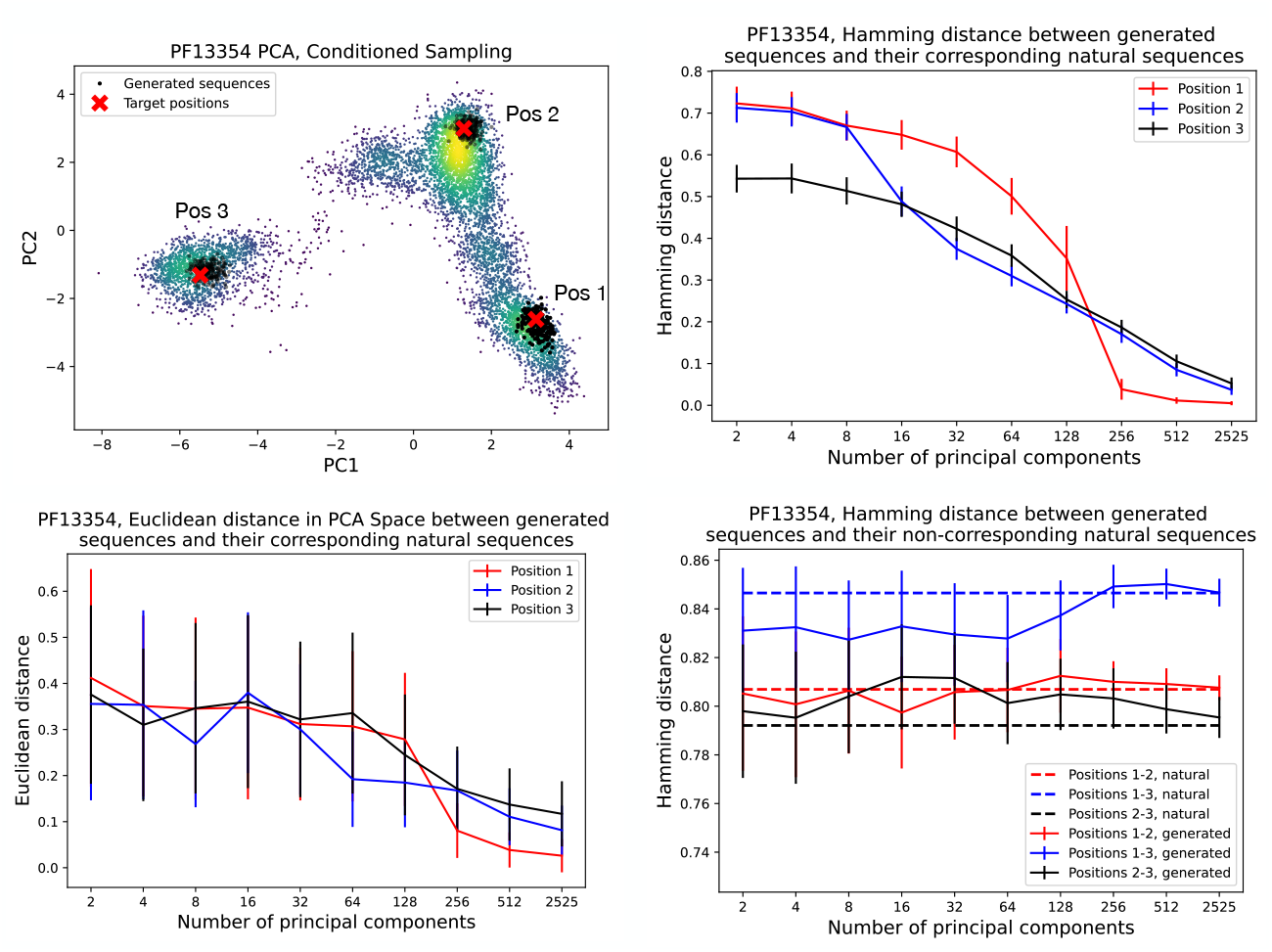
Statistical analysis of the generativity conditioned on three different positions on the PC space as a function of the number of principal components learned during training. **Top left**: red crosses represent the three positions chosen on different islands of the PC projection to study the different conditioned sampling. The clouds of black dots represent the generated sequences around the target positions. **Bottom left**: Euclidean distance in the first two PC plane between the target positions and the generated sequences conditioned on those positions. **Top right**: Hamming distance between the target positions and the generated sequences conditioned on those positions. **Bottom right**: Hamming distance between the generated sequences and the non-corresponding target positions.

Using structure prediction models such as AlphaFold 3 (AF) [23] or ESMFold (ESM) [19], we can evaluate the folding of generated sequences as a function of the number of principal components. Starting from two wild-type sequences with experimentally determined crystal structures in the Protein Data Bank (PDB) [27], we fold the generated sequences using ESM and compare their predicted structures to the experimental ones via root-mean-square deviation (RMSD) of atomic positions. However, this analysis is inherently limited by the accuracy of ESM or AF in distinguishing structurally diverse members within the same protein family. In some families, reliable evaluation is not possible, as both ESM and AF fail to capture distinct foldings among naturally aligned but structurally divergent sequences. Panels in Figure 4 illustrate two contrasting cases for protein families PF00014 and PF00076. In the first case (PF00014), the RMSD between the predicted structure of the two wild-type sequences (blue dotted line) does not match the RMSD between their experimental structures (red dotted line). As a result, the sequences generated with FeatureDCA from either wild type are both folded into the structure of wild type 1. In contrast, for PF00076, the structural predictions of ESM closely match the experimental structures, effectively distinguishing between the two conformations. In this case, the sequences generated with FeatureDCA fold into three-dimensional structures that are closely aligned with their respective wild types.

**Figure 4:**
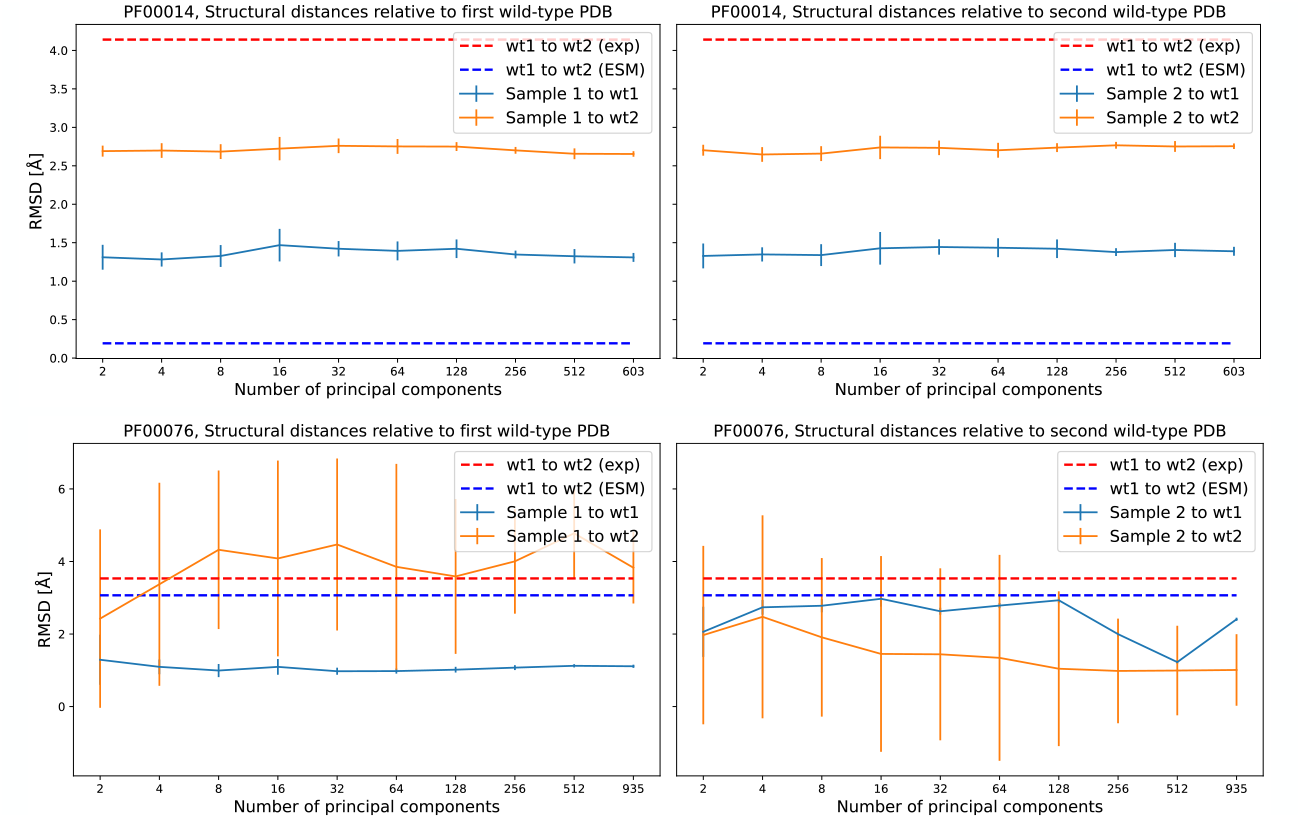
Structural similarity of generated sequences to wild-type structures in protein families PF00014 and PF00076. Each panel shows the RMSD (Å) between the predicted structures of generated sequences and their respective wild-type references, plotted against the number of principal components used for learning and sequence generation. **Top**: PF00014; **Bottom**: PF00076. **Left**: RMSD relative to the first wild-type (wt1) structure; **Right**: RMSD relative to the second wild-type (wt2) structure. Dashed lines indicate the RMSD between the two wild types using experimental structures (red) and ESM predictions (blue). Solid lines represent average RMSDs of generated samples to wt1 (blue) and to wt2 (orange), with error bars showing standard deviation. This analysis highlights the structural consistency of sampled sequences relative to the wild types and the capacity of ESM to recover alternative foldings, which appears limited in PF00014 but is more variable in PF00076.

#### The case of RR homodimers

The generative capabilities of FeatureDCA can be evaluated on a protein domain characterized by both a structurally diverse landscape and a rich sequence repertoire, ensuring that accurate folding predictions can be obtained across its variants using models such as AlphaFold (AF) or ESMFold (ESM). A prototypical example is the bacterial Response Regulator (RR) family (Pfam ID PF00072), which functions as part of a two-component signal transduction system. In this system, a sensor histidine kinase (HK) detects environmental cues and undergoes autophosphorylation, subsequently transferring the phosphate group to its RR partner. Phosphorylation activates the RR, typically inducing a conformational change that alters its function, most commonly by pro-moting DNA binding and transcriptional regulation. RRs exhibit architectural diversity depending on their DNA-binding domains, which in turn influence their dimerization mechanisms [28, 29, 30]. Among the PF00072 sequences, we focus on three well-characterized subclasses that form distinct homodimers upon phosphorylation. These dimerization modes correspond to distinct domain architectures, each comprising the conserved receiver domain PF00072 combined with one of three DNA-binding domains: Trans Reg C (PF00486), GerE (PF00196), or LytTR (PF04397). Figure 5 shows representative homodimer structures from the three RR subclasses (A–C) and their pairwise structural alignments (D–F), performed using PyMOL’s `align` function [31], which reports the root-mean-square deviations (RMSDs) between matched atoms. The PDB IDs corresponding to the Trans Reg C, LytTR, and GerE classes are 1NXS, 4CBV, and 4ZMS, respectively. The RMSDs for the three pairwise alignments are: 8.6 Å between Trans Reg C and LytTR (D), 16.1 Å between Trans Reg C and GerE (E), and 18.7 Å between LytTR and GerE (F).

**Figure 5:**
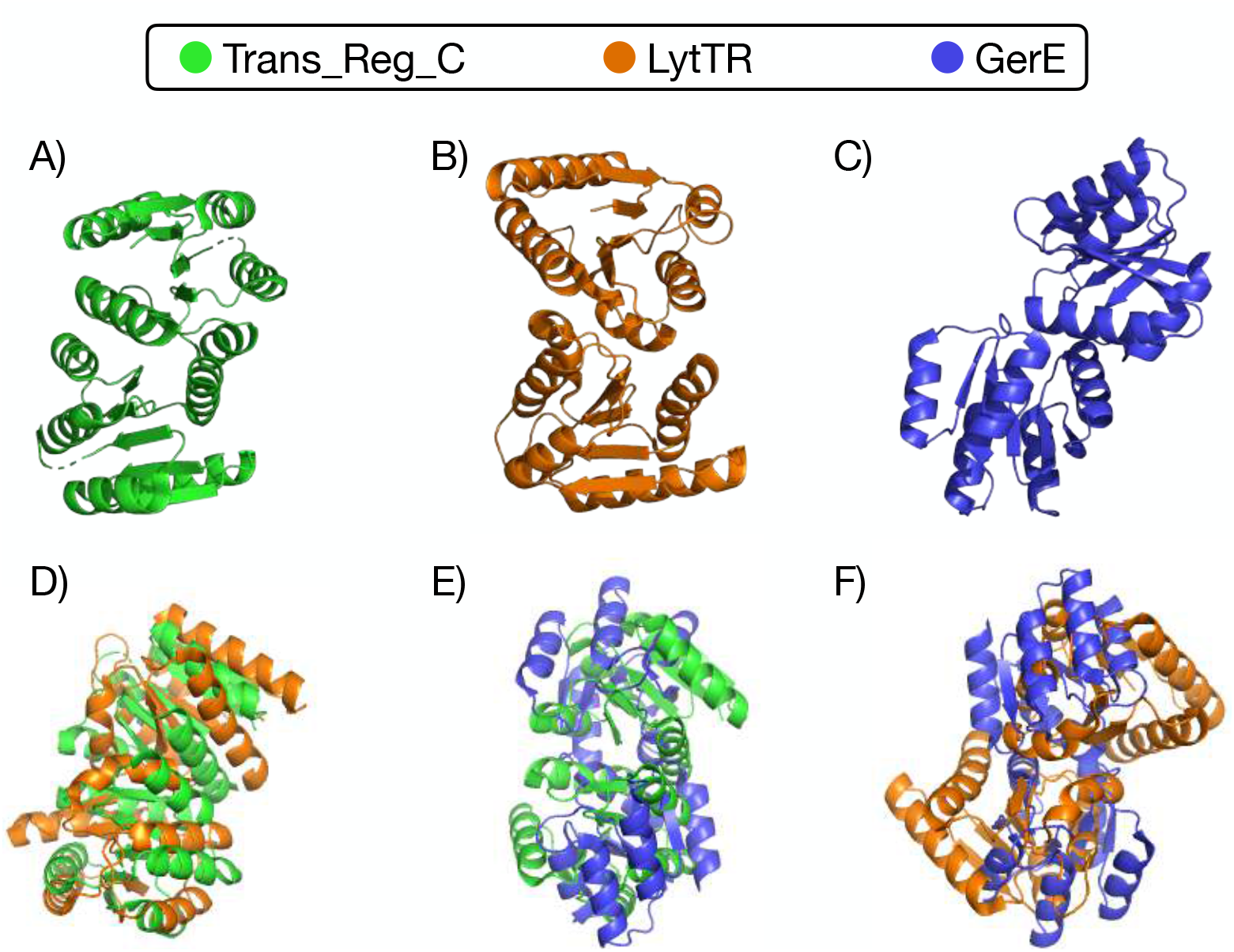
Structural comparison of three Response Regulator (RR) subclasses with distinct dimerization modes. **A–C)** Experimentally determined homodimer structures of representative RR proteins containing the DNA-binding domains Trans Reg C (A, PDB ID: 1NXS), LytTR (B, PDB ID: 4CBV), and GerE (C, PDB ID: 4ZMS). **D–F)** Pairwise structural alignments between these dimers, performed using PyMOL’s align function. The root-mean-square deviations (RMSDs) between the dimers quantify the extent of structural divergence: 8.6 Å between Trans Reg C and LytTR (D), 16.1 Å between Trans Reg C and GerE (E), and 18.7 Å between LytTR and GerE (F). These values highlight the substantial differences in quaternary structure and dimerization geometry across the three RR subclasses, despite all sharing the same conserved receiver domain fold.

To evaluate the generative capabilities of FeatureDCA, which learns from and generates aligned sequences, it is essential to verify that AF or ESM can accurately fold natural sequences from the alignment into the correct homodimer classes observed in experimental structures. Among the 1.7 million UniProt sequences assigned to PF00072, we identified 160, 585 with an architecture compatible with the Trans Reg C class, 69, 401 with GerE, and 34, 699 with LytTR, for a total of 264, 685 non-redundant sequences. Based on these, we constructed a multiple sequence alignment (MSA) of 118 positions, longer than the 111-position Hidden Markov Model (HMM) profile available in Pfam (now integrated into InterPro)[32, 33], ensuring coverage of the relevant structural features. When folded with AlphaFold 3, the aligned natural sequences adopt structures highly consistent with their experimental counterparts: the root-mean-square deviations between predicted and crystal structures are 0.35 Å for 1NXS (Trans Reg C), 1.1 Å for 4CBV (LytTR), and 1.3 Å for 4ZMS (GerE). These values fall within or below the resolution range of the experimental structures (1.5–4 Å), confirming the structural compatibility of the alignment for downstream generative modeling. An interesting feature of the dataset is that the three subsets corresponding to the homodimer classes form well-separated clusters when projected onto the first two principal components. This clear separation in sequence space, as shown in Figure 6, supports the idea that FeatureDCA can learn to condition sequence generation on structural class.

**Figure 6:**
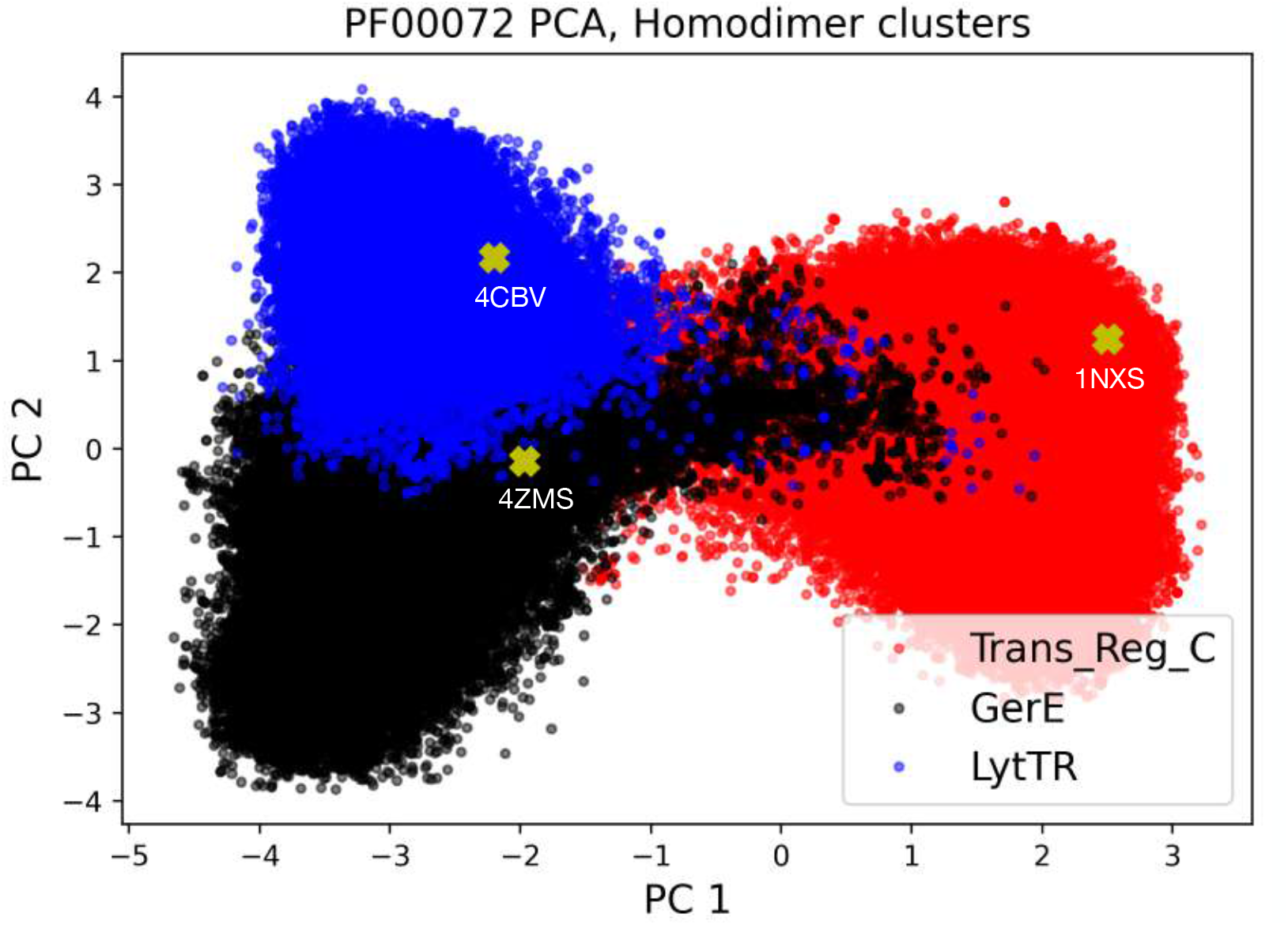
PCA projection of aligned RR sequences reveals distinct structural clusters. Projection of the multiple sequence alignment of PF00072 sequences onto the first two principal components. The three main clusters correspond to sequences containing the Trans Reg C (red dots), LytTR (blue), and GerE (black) DNA-binding domains, associated with distinct homodimerization classes. Yellow crosses mark the positions of the experimental PDB structures 1NXS (Trans Reg C), 4CBV (LytTR), and 4ZMS (GerE), each falling within its respective cluster. This separation supports the idea that structural class is encoded in sequence space and can be learned by FeatureDCA.

To assess this claim, we performed a targeted sampling experiment. For each structural class, we extracted the principal component (PC) coordinates of the sequences corresponding to the experimental structures (1NXS, 4ZMS, or 4CBV) and used them as conditioning inputs to generate new sequences. These generated sequences were then folded using AlphaFold 3 and compared, via RMSD of the aligned atomic positions, to all three experimental wild-type structures. The top-right, top-left, and bottom-left panels in Figure 7 show the results of this experiment. Each panel displays the average RMSD between the predicted structures of generated sequences and all three wild-type references, plotted as a function of the number of principal components used during model training and generation. The RMSD to the structure of the same class used for conditioning is expected to decrease with increasing number of PCs, while the RMSDs to the other two reference structures tend to approach the experimentally measured distances between the corresponding wild-type structures (dotted lines).

**Figure 7:**
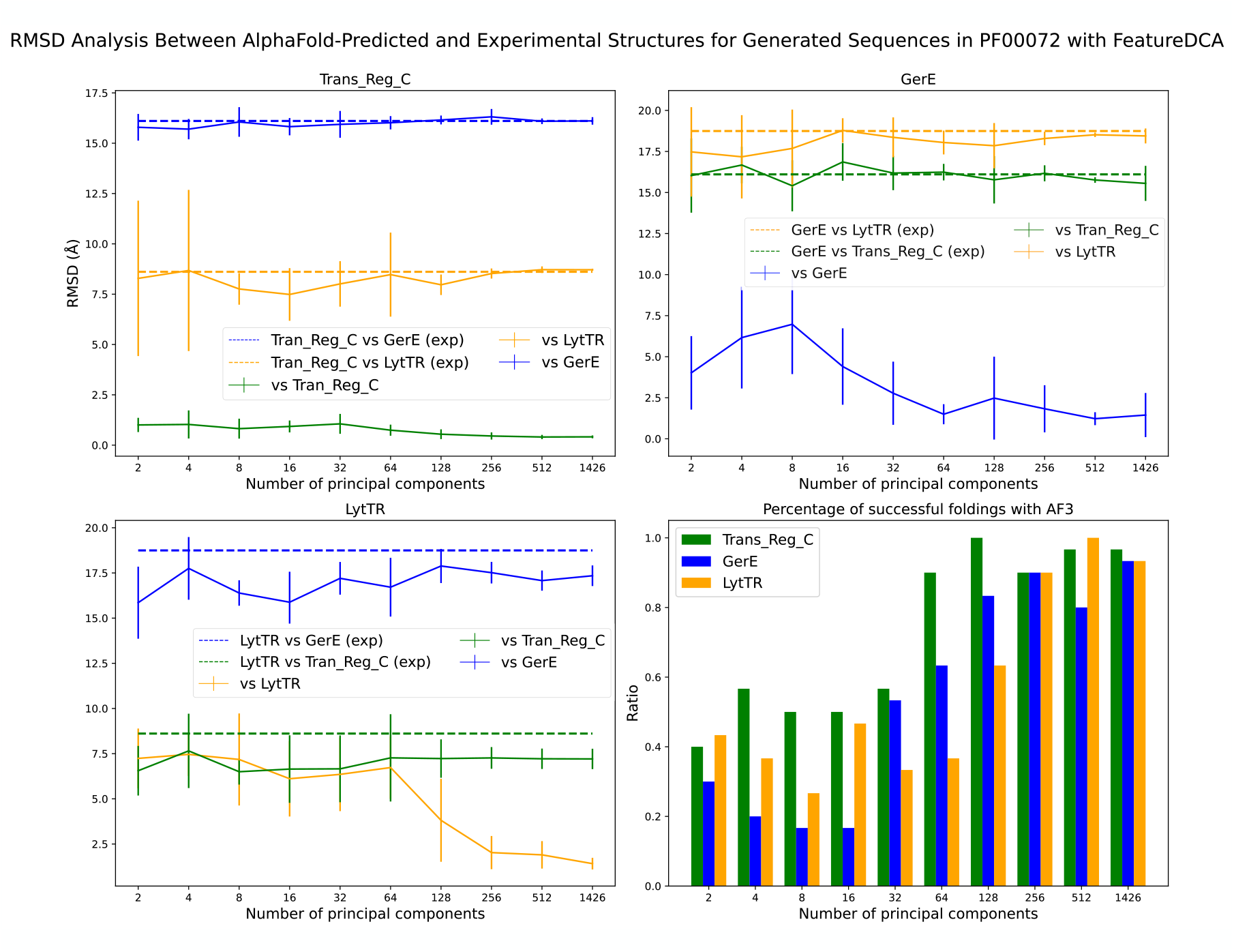
Structural consistency and class specificity of FeatureDCA-generated sequences for the PF00072 protein family. The **top-left, top-right**, and **bottom-left** panels show the root-mean-square deviation (RMSD) between predicted structures of generated sequences from one of the three RR structural classes, Trans Reg C (PDB ID: 1NXS), GerE (PDB ID: 4ZMS), and LytTR (PDB ID: 4CBV), respectively, and the three experimental wild-type structures. For each class, sequences were generated by conditioning FeatureDCA on the principal component (PC) coordinates of the sequences corresponding to the experimental PDB structures, and the predicted structures of these generated sequences were compared to all three references using AlphaFold 3. Solid lines represent the average RMSD as a function of the number of PCs used during training and generation. In each plot, the curve corresponding to the reference structure of the same class (e.g., 1NXS for Trans Reg C, green curve in the top right panel) is expected to show the lowest RMSD, while the other two curves converge toward the RMSDs between the experimental structures (shown as dotted lines), reflecting the divergence across classes. The **bottom-right** panel shows the fraction of generated sequences whose predicted structures are within 10 Å RMSD of the corresponding class-specific wild-type structure, indicating successful fold recovery. This analysis demonstrates that FeatureDCA not only generates structurally plausible sequences, but also preserves class-specific structural identity as a function of PC-based conditioning.

The trend shown in Figure 7 confirms that FeatureDCA can effectively guide generation toward the intended structural class, provided that sufficient principal component information is used during training. However, there are clear differences in class-specific performance. The Trans Reg C class (1NXS), which is the most represented in the dataset, is consistently reproduced with the lowest RMSDs, indicating that FeatureDCA learns its statistical and structural features more effectively. In contrast, the LytTR class (4CBV) presents a greater challenge: for low-dimensional conditioning, its generated sequences often adopt structures more similar to Trans Reg C than to LytTR itself, suggesting confusion between classes. Only when the number of PCs becomes sufficiently large does the model begin to distinguish LytTR with sufficient resolution. These observations suggest that both training-set representation and feature dimensionality play key roles in enabling class-specific generative control. Finally, the bottom-right panel in Figure 7 quantifies the success rate of folding the generated sequences with AlphaFold. A fold is considered successful if its predicted structure lies within a threshold RMSD to the corresponding target structure; here, the threshold is set to 10 Å. Alternatively, one could assess folding quality using the predicted template modeling (pTM) score returned by AlphaFold, and we find that the same qualitative conclusions hold: sequences conditioned on well-sampled structural classes are more likely to produce high-confidence, accurate folds.

The results confirm that FeatureDCA, when appropriately conditioned, can generate structurally plausible sequences that reflect subclass-specific geometries, but also highlight the limits of lowdimensional conditioning in sparsely represented regions of sequence space.

### Predicting mutational effects via in-silico deep mutational scanning

An important test for sequence-based models is their ability to capture the functional impact of mutations. In particular, in-silico deep mutational scanning (DMS) is a key application of statistical models trained on MSAs, offering the ability to predict the fitness effects of single-residue mutations by computing the log-likelihood difference between wild-type and mutant sequences:

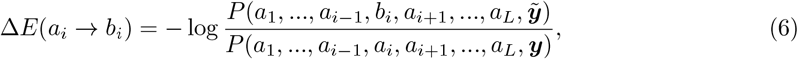

where *P* (***a, y***) represents the joint probability distribution of the autoregressive model defined by Equations 1, 4, while ***y*** and 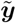 are the PC projections of the wild-type and mutant sequence, respectively.

Following the approach of Trinquier et al. [5], we benchmarked FeatureDCA against ArDCA on the well-studied Beta-lactamase family (PF13354), for which experimental DMS data are available [34, 35, 7, 36].

We evaluated the Spearman rank correlation between the predicted and experimental mutational effects. As shown in SI Section 7, FeatureDCA achieves comparable performance to ArDCA for small values of the conditioning dimension *d*. Interestingly, as the number of principal components used during training increases, we observe a non-monotonic behavior in predictive performance. Specifically, the correlation initially decreases, suggesting that conditioning on an excessive number of PCs may overfit or dilute mutational signals. However, at larger values of *d*, the performance begins to recover, indicating that the model eventually re-stabilizes and incorporates additional global variation effectively.

These results demonstrate that FeatureDCA not only supports conditional generative modeling but also remains competitive in mutational effect prediction, with performance that can be tuned via the dimensionality of the conditioning space.

## Discussion

In this work, we presented FeatureDCA, a feature-conditioned extension of the Direct Coupling Analysis (DCA) framework, designed for controllable generation of protein sequences within a given protein family. By integrating biologically relevant, low-dimensional features, specifically the principal components (PCs) derived from multiple sequence alignments (MSAs), FeatureDCA enables users to direct sequence generation toward targeted regions of sequence space, all while preserving the statistical and structural characteristics of natural proteins used for training.

Through extensive simulations, we showed that FeatureDCA accurately reproduces key aspects of protein families, including coevolutionary statistics, principal component distributions, and structural properties. Notably, the model generates novel sequences that remain at a substantial Hamming distance from training examples, supporting diversity, an essential criterion in protein design.

When compared to established models such as bmDCA and ArDCA, FeatureDCA delivers competitive or superior generative performance and introduces a crucial new capability: directional control over the generated sequence distribution, as illustrated in Figure 2.

We also showed that FeatureDCA can perform localized generation around user-defined points in feature space, allowing the model to sample sequences that approximate specific wild-type sequences both statistically and structurally, see Figures 3, 4. In particular, the case study on the Response Regulator (RR) family (PF00072) highlighted FeatureDCA’s capacity to navigate intra-family structural diversity, similarly to what is shown in [21]. By conditioning on PC coordinates of known RR subtypes, shown in Figures 5 and 6, the model was able to selectively generate sequences that folded into class-specific dimerization geometries, demonstrating its potential for targeted generative design. In addition, we showed that FeatureDCA retains predictive power on experimental deep mutational scanning data, further confirming its ability to capture local functional constraints in sequence space.

However, limitations remain. First, the conditioning space is currently restricted to principal components that, while biologically meaningful, are not explicitly linked to function or structure. Extending FeatureDCA to condition on experimentally or computationally derived features, such as binding specificity, fitness, or structural class, would broaden its applicability without changing the architecture of the outlined model. Second, generative performance in sparsely populated regions of feature space, such as the LytTR subclass for the case of RR domains, is limited, reflecting the challenge of learning from underrepresented sequence types. More balanced or augmented training sets may help alleviate this problem.

A key strength of FeatureDCA lies in its flexibility. Unlike standard Potts-based models that rely on global sampling schemes, the autoregressive implementation used to solve the model allows for efficient training and generation by factorizing the sequence distribution into a chain of conditionals. This not only makes FeatureDCA computationally comparable to ArDCA but also allows the sampling process to be explicitly guided by external features beyond local amino acid frequencies. As demonstrated in this work, such conditioning enables both global and localized control over the generative process, opening the door to a broad range of applications in statistical modeling and protein design.

In summary, FeatureDCA bridges the gap between interpretable statistical models and flexible, feature-guided protein design. It provides a robust and efficient framework for controllable sequence generation grounded in evolutionary data, with the potential to guide functional exploration, structural modeling, and synthetic biology applications.

## Data and Code Availability

All datasets used in this work are publicly available at: https://github.com/francescocaredda/FeatureDCAData. The full Julia implementation of `FeatureDCA`, including example Jupyter notebooks for reproducibility and application, can be found at: https://github.com/francescocaredda/FeatureDCA.jl. These repositories provide all the necessary resources to replicate the experiments and analyses described in this study.

## Acknowledgments

We are deeply grateful to Martin Weigt and Leonardo Di Bari for many interesting discussions on addressable sequence generation. AP and FC acknowledge financial support from the project “Explainable Models for Protein Design”, funded by the MIUR Progetti di Ricerca di Rilevante Interesse Nazionale (PRIN) Bando 2022 - grant 2022TE5B7X. We also acknowledge “Centro Nazionale di Ricerca in High-Performance Computing, Big Data and Quantum Computing” (ICSC). This study was carried out within the “FAIR - Future Artificial Intelligence Research” project, and received funding from the European Union NextGenerationEU (Piano Nazionale di Ripresa e Resilienza (PNRR)–Missione 4 Componente 2, Investimento Grants No. 1.3–D.D. 1555 11/10/2022, and No. PE00000013). This paper reflects only the authors’ views and opinions, neither the European Union nor the European Commission can be considered responsible for them.

## Supplementary Material

### A Direct Coupling Analysis Details

#### Summary Statistics

A Multiple Sequence Alignment 𝒟 composed of *M* sequences of length *L* can be used to extract summary statistics representing the protein family. In particular, single- and pair-wise frequency counts of the amino acids in the alignment are defined as:

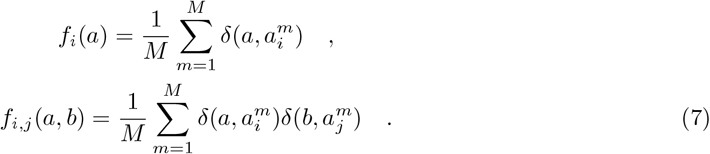

The application of a maximum entropy principle to the frequency counts returns the Potts model at the base of every DCA method. Higher statistics would produce other terms in the Hamiltonian which could not be inferred due to the relatively small size of the datasets at hand [24, 25].

A good generative model should be able to reproduce the summary statistics and higher statistics derived from it, as in the case of the 2-site connected correlation:

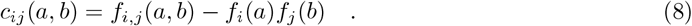

#### Sequence re-weighting and effective number of sequences

Sequences in a natural MSA are not truly independently distributed. Indeed, a certain degree of homology arises due to the phylogenetic relationships between different sequences, bringing about probabilistic biases towards specific sets of sequences. To avoid the over-sampling of similar sequences and the relative under-representation of other sequences, it is necessary to introduce a reweighting scheme. For each sequence **a**^*i*^, a weight is defined as the inverse of the number of sequences similar to it:

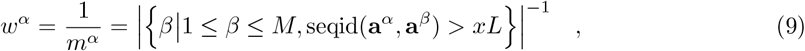

where *x* is a similarity threshold that we set equal to 0.9. The effective number of sequences (effective depth) of the MSA *M*_eff_ is then defined as the sum of each weight:

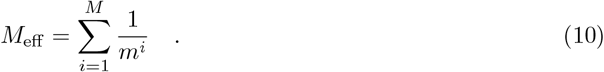

Using this re-weighting scheme, the frequency counts can be written as:

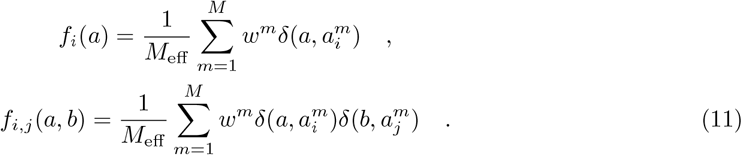

Depending on the effective depth of an MSA, a given DCA method can be more or less efficient, with different methods presenting different precision thresholds [25].

### B Mathematical foundation of the feature-conditioned au-toregressive model

Let us assume that we have an MSA of *M* one-hot encoded sequences of length *L* over an alphabet of *q* symbols (*q* = 21 for proteins). We can extract the rotation matrix *U* ∈ ℝ^*d×Lq*^, such that the application *U* **a** = **y** is the projection of an amino-acid sequence **a** in the *d*-dimensional space spanned by the eigenvectors associated to the largest *d* eigenvalues of covariance matrix *C*^*emp*^. Let us define the sequence

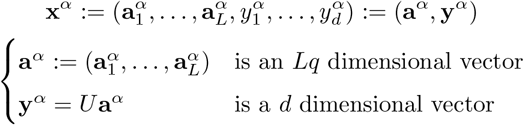

where **a**_*i*_ is the *q*-dimensional one-hot encoded representation of amino-acid *a*_*i*_. We aim at learning the joint probability distribution *P* (**a, y**) from our data. We note that

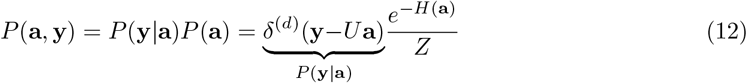

where: (i) we assume in the last step that the prior distribution of **y** is uniform, 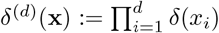 is the *d*-dimensional Kronecker delta, and (ii) we model *P* (**a**) as a Potts model defined by the energy function *H*(**a**). The joint probability distribution *P* (**a, y**) is amenable to autoregressive representation:

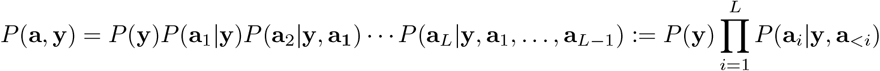

where **a**_*<i*_ := {*a*_1_, …, *a*_*i−*1_} with the convention that **a**_*<*1_ is the empty set.

#### Gaussian Approximation

To proceed analytically, we make use of a Gaussian approximation for *P* (**a**) = exp(−*H*)*/Z*. Also, we relax the one-hot-encoded nature of the variables **a**_*i*_ to real values. In this case, we have that

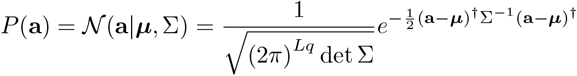

Eq. 12 becomes:

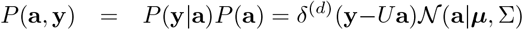

We can compute the marginal *P* (**y**) which is a Gaussian

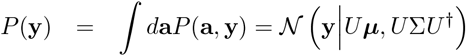

#### Computation of the marginal *P* (***y***)

Let us consider the case *P* (**y**|**a**) = *δ*^(*d*)^(**y**−*U* **a**). We have that

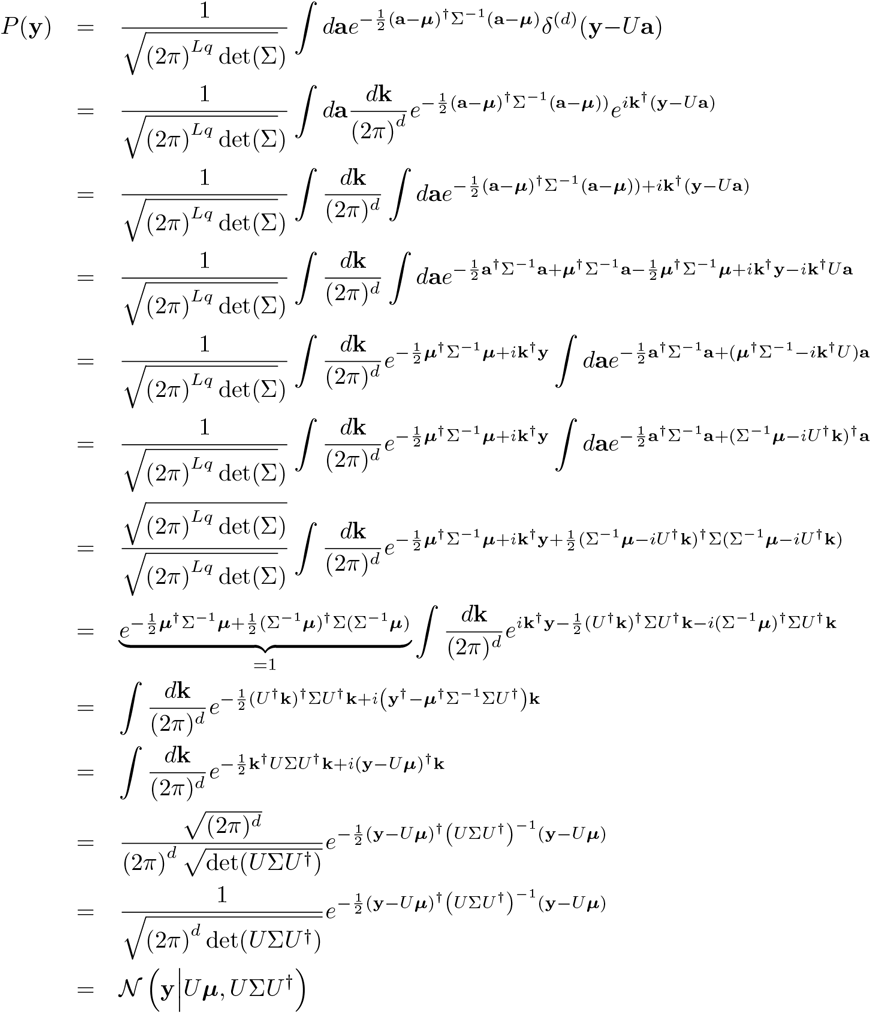

Alternatively, we could have computed the same result by observing that *P* (**y**) is Gaussian and its first two moments can be computed as:

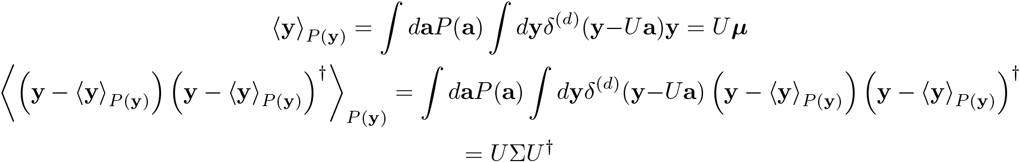

In the following, we will use this technique extensively.

#### Computation of *P* (a|y) and its marginals

So far, we can write the conditioned distribution *P* (**a**|**y**) as

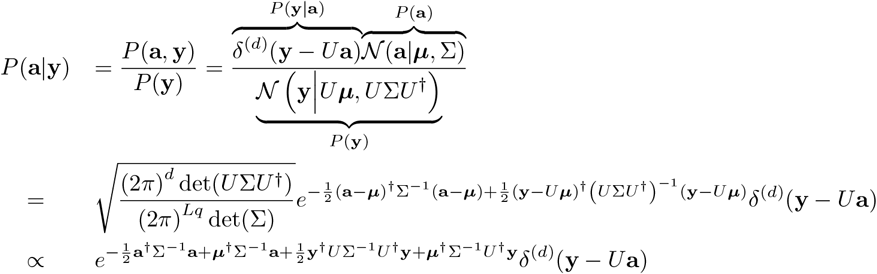

so that

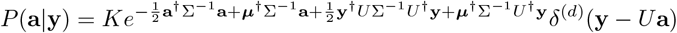

where *K* is a normalization constant that does not affect the following computations.

##### First marginal

The first marginal of the conditioned distribution *P* (**a**|**y**) can be computed as

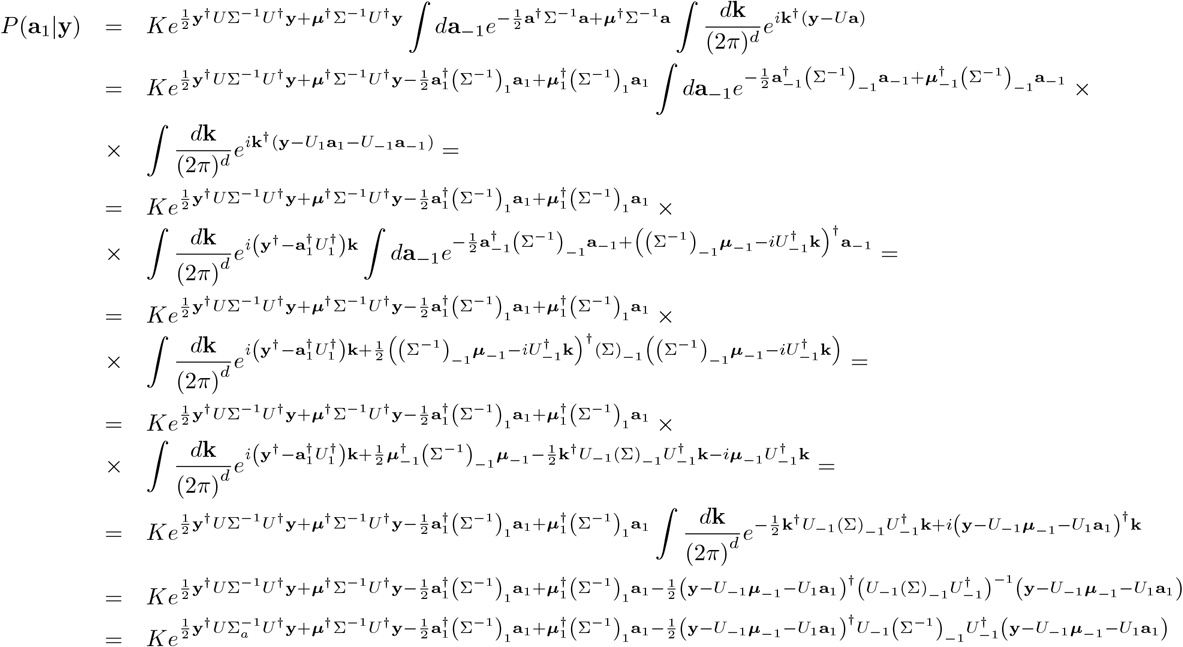

The marginal distribution *P* (***a***_1_|***y***) remains a Gaussian distribution that can be fully characterized by its mean and variance. The mean can be computed as the maximum of the argument of the exponential, while the variance as the opposite of the inverse of its second derivative with respect to the variable ***a***_1_ :

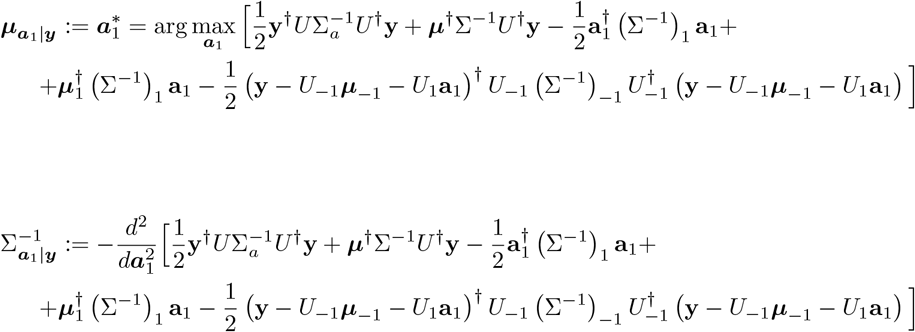

Recalling the identities:

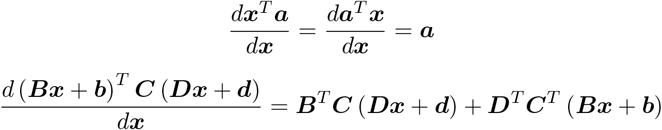

we have that the mean is given by:

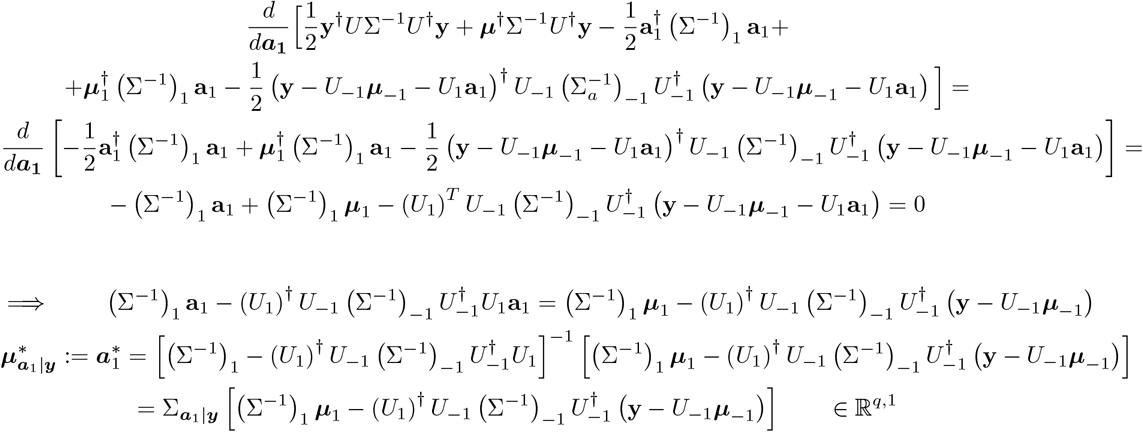

while the variance is:

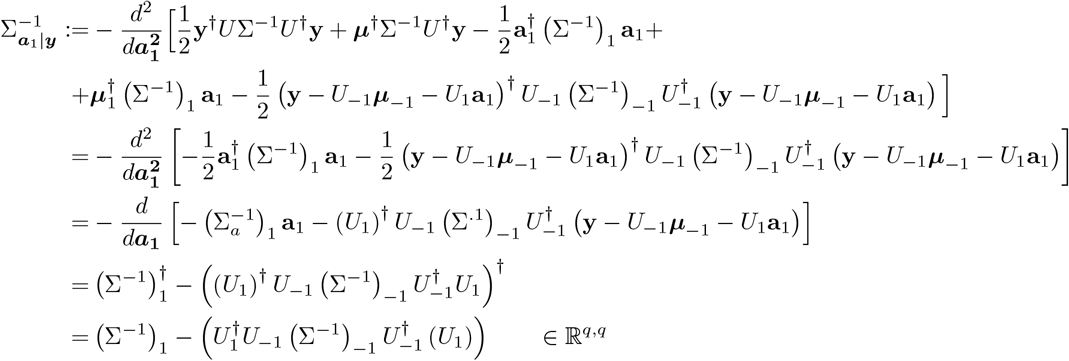

So the marginal can be written as

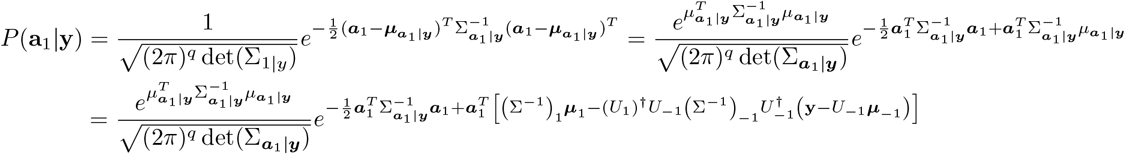

In this context the notation (Σ^*−*1^ ∈ ℝ^*q,q*^) refers to the first block of matrix Σ^*−*1^, while Σ^*−*1^ ∈ ℝ^*q*(*L−*1),*q*(*L−*1)^ refers to the sub-matrix defined by removing the blocks relative to the first amino-acid. The same goes for *U*_1_ ∈ ℝ^*d,q*^ and *U*_*−*1_ ∈ ℝ^*d,q*(*L−*1)^.

##### *k*-th marginal

Generalizing the previous computation, we can compute the *k*-th marginal of the distribution *P* (***a***_*k*_|***a***_*<k*_, ***y***) conditioned to all previous positions and to PCA components ***y***. Using Bayes’ rule we can write:

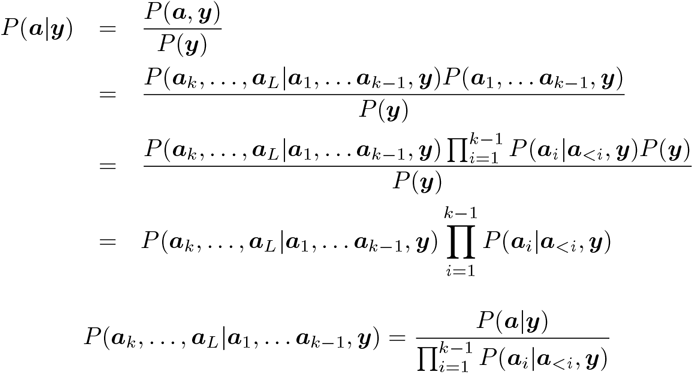

From this, we can define the marginal as:

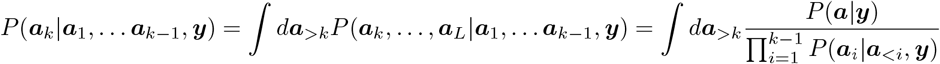

The actual computation is quite straightforward

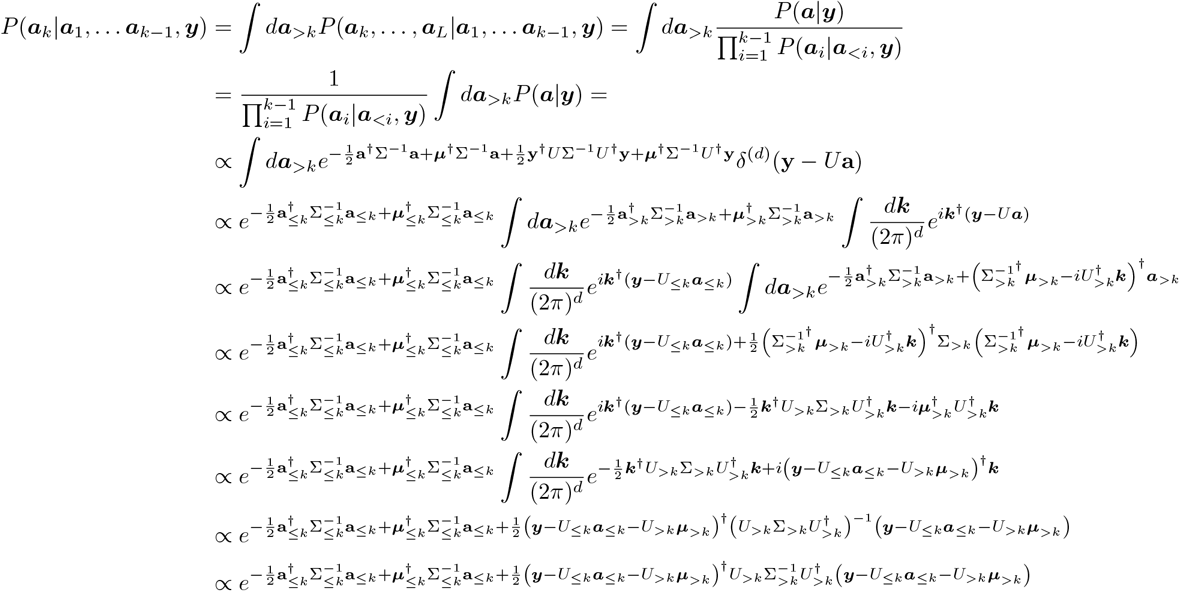

Now it is essential to isolate the terms depending on ***a***_*k*_ alone in order to being able to compute the first and second derivatives to define mean and variance of the gaussian. In the following, indices *i, j* refer to block elements of the matrices they are applied to, while the term *constant* is used for terms that do not explicitely depend on **a**_*k*_. Let’s analyze each term in the exponential of the marginal distribution:

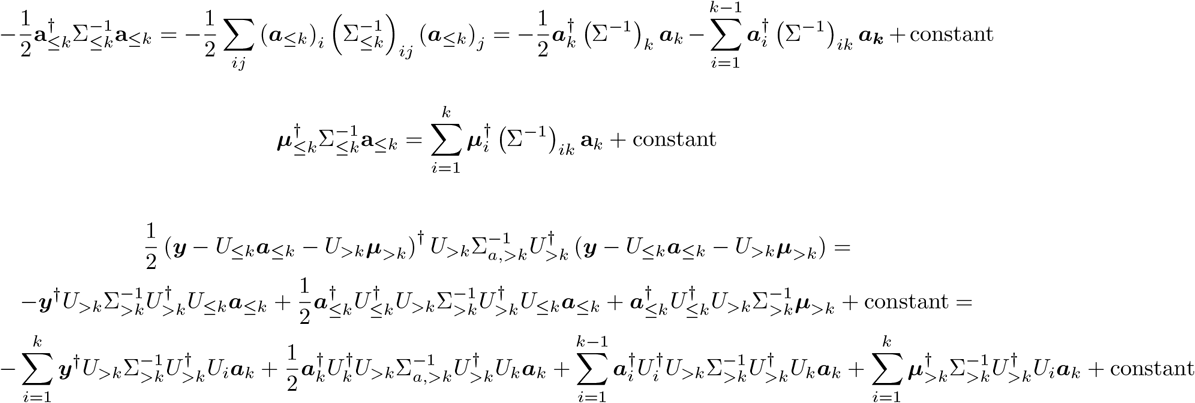

Computing the first derivative with respect to ***a***_*k*_ of the argument of the exponential and setting it equal to zero, we get:

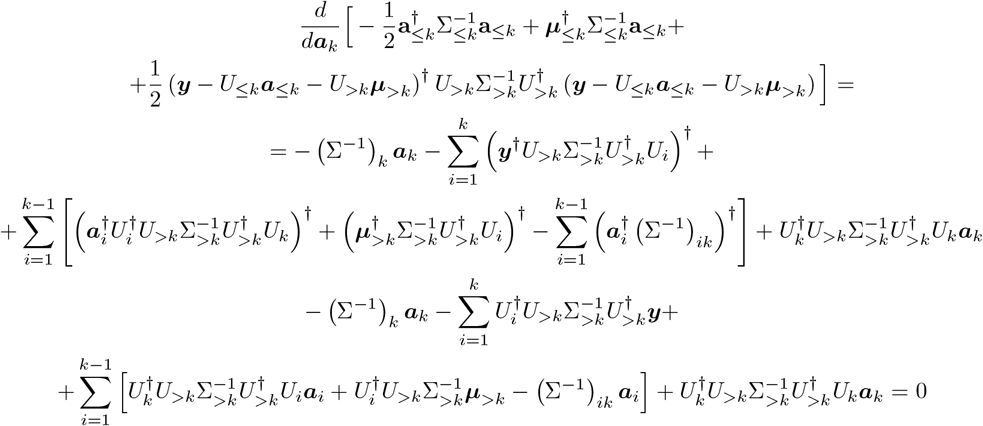

which gives a mean 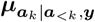 given by:

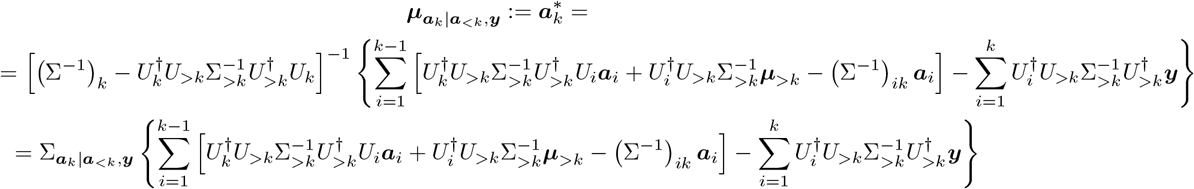

The second derivative returns the inverse of the covariance matrix

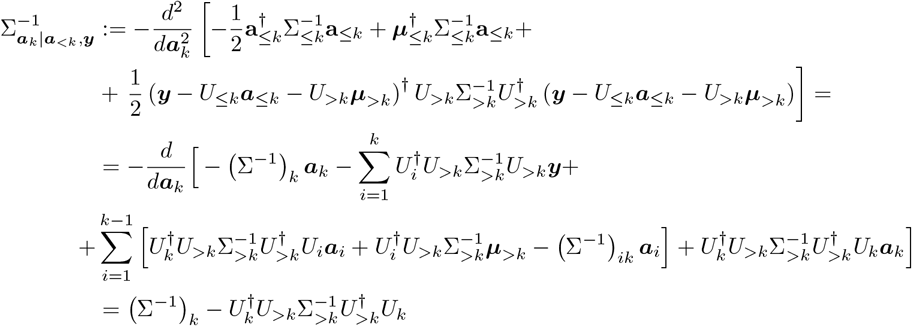

Having characterized the *k*-th marginal, we have a solid foundation to define the *k*-th term in the autoregressive decomposition of the full joint distribution *P* (**a, y**):

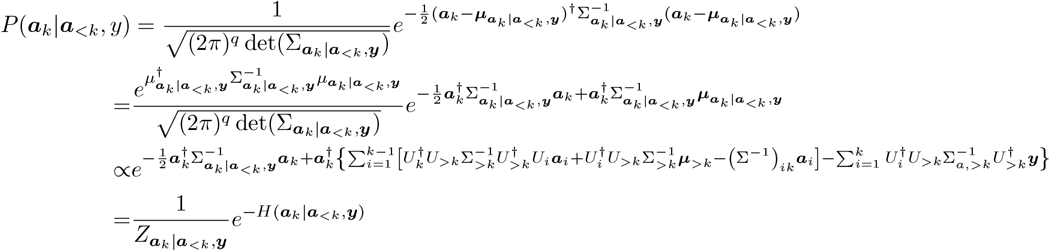

where the energy function *H*(**a**_*k*_|**a**_*<k*_, **y**) can be written as:

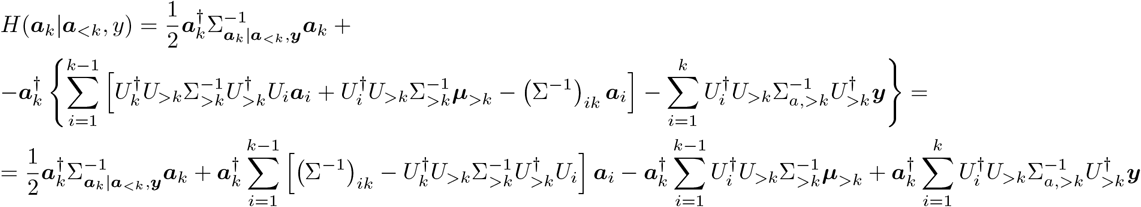

### Discrete model

Now that we have a principled formulation of the energy function of the ***k***−th term in the autore-gressive decomposition, we can restore the discrete nature of the amino-acid data and produce an ansatz for a discrete version of the model in its autoregressive implementation. Let’s consider each of the three contributions in the energy function:

1. Quadratic term in the amino-acid space

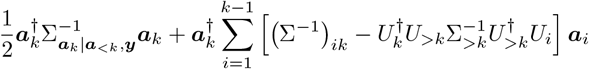 This can be thought of as an interaction term:

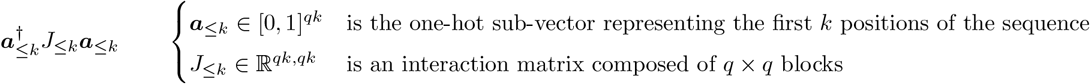 Using a *one-cold* representation where ***a*** ∈ {0, 1, 2, …, *q*}^*L*^, the same can be written as

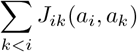
2. Linear term in the the amino-acid space

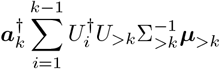 This can be easily interpreted as a local field. In a one-hot representation this is 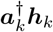 with ***h***_*k*_ ∈ ℝ^*q*^, while in a *one-cold* representation this is

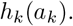
3. Cross interaction term between the amino-acid space and the principal component space:

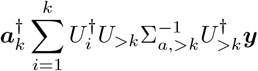

This can be written as

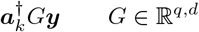

where *G* acts as an embedding mapping from the amino-acid space to the principal component space. In a *one-cold* representation, this is

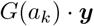

The full one-cold energy function thus reads:

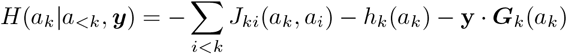

and the probability distribution of the *k*-th term is

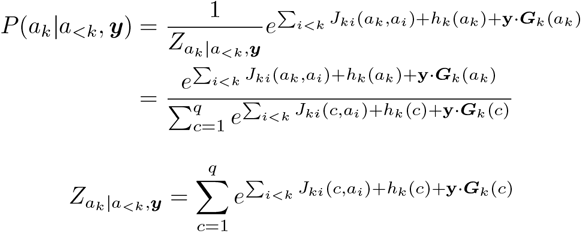

Given a MSA 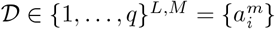 concatenated to a matrix 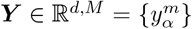 representing the principal components of each sequence in 𝒟, the likelihood of the model can be written as

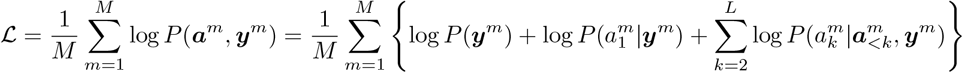

The first term of the likelihood can be thought of as an empirical frequency of the sequences in the PC space. Since it does not depend on the learning parameters of the model, it can be discarded.

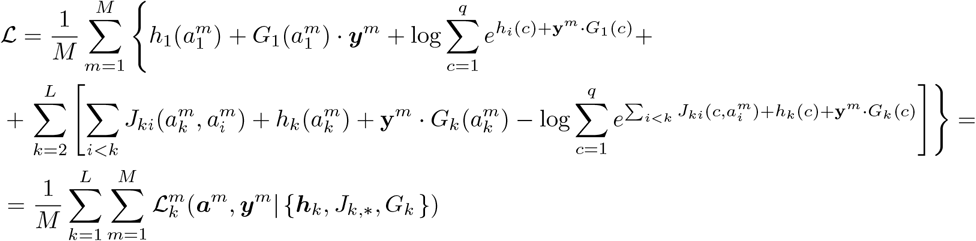

Since the parameters relative to each site of the sequence are factorized, the full likelihood can be optimized in parallel. So, each single optimization problem is defined by the following objective function:

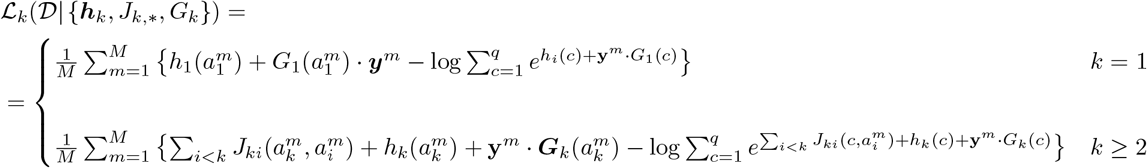

### C Implementation Details

The model architecture and all computational experiments were implemented in the Julia programming language. The full source code, along with documentation and scripts to reproduce the results, is available in a public GitHub repository https://github.com/francescocaredda/FeatureDCA.jl.

Model training proceeds by minimizing the negative log-likelihood of the conditional distribution at each position *k*, corresponding to the autoregressive factorization described in Eq. (1) of the main text. The objective function was optimized using deterministic gradient descent with the `LD LBFGS` algorithm [37] provided by the `NLopt.jl` package [38]. Optimization was performed with a convergence tolerance of 10^*−*5^. We applied *L*_2_ regularization to three model components: the local fields *h*_*i*_(*a*), the interaction tensors *J*_*ij*_(*a, b*), and the feature embedding matrix *G*. A single regularization coefficient of *λ* = 10^*−*4^ was used across all terms.

Principal component analysis (PCA) was conducted using the `MultivariateStats.jl` package. Multiple sequence alignments (MSAs) were one-hot encoded and centered prior to PCA. The top *d* principal components were retained and used as conditioning inputs throughout training and generation.

Training and sampling can be parallelized across sequences or positional blocks due to the factorized structure of the model. Sinkhorn divergences were computed using a custom Julia implementation of the Sinkhorn–Knopp algorithm [26] in the `OptimalTransport.jl` package.

### D Data Processing and MSA Construction

The analyses presented in the *Generativity* section of the main text were carried out using the multiple sequence alignments (MSAs) available at https://github.com/pagnani/ArDCAData, which were originally curated for the benchmarking of ArDCA [5].

For the analysis of the Response Regulator (RR) family (Pfam ID PF00072), we constructed a custom MSA to capture the structural diversity across its known functional subclasses. We first retrieved from UniProt all sequences exhibiting a two-domain architecture composed of the conserved receiver domain PF00072 in combination with one of the three DNA-binding domains that define the major RR subclasses: PF00486 (Trans Reg C), PF04397 (LytTR), and PF00196 (GerE). This yielded a total of 160,585 non-redundant sequences. To ensure structural relevance and compatibility with available experimental structures, we used the `jackhmmer` tool to identify the 200 sequences in UniProt most similar to the RR representatives in the Protein Data Bank: 1NXS (Trans Reg C), 4CBV (LytTR), and 4ZMS (GerE). These sequences were used to construct a profile Hidden Markov Model (HMM) using the `hmmbuild` tool from the HMMER suite [39]. The resulting HMM spanned 118 aligned positions and was used as a reference for alignment. All 160,585 RR sequences were then aligned to the profile HMM using `hmmalign.` Insertions relative to the HMM model were removed to ensure consistency in positional indexing, resulting in a clean, fixed-length MSA. This alignment was used to train FeatureDCA for the generation of class-specific RR sequences as described in the section *The case of RR homodimers* of the main text. Data relative to the study of RR homodimers is available at https://github.com/francescocaredda/FeatureDCAData

### E Wasserstein distance and Sinkhorn divergence

Comparing the PCA projections of natural and generated sequences is a common step in evaluating generative models for aligned protein families. This comparison is often performed qualitatively, by inspecting two-dimensional projections and assessing whether the generated sequences reproduce the overall geometry, clustering, and density of the natural distribution. To complement this visual inspection with a rigorous, quantitative metric, we use the Wasserstein distance, a concept originating from Optimal Transport theory, which provides a means to measure the distance between two probability distributions over a geometric space.

Consider two distributions of points **X** = {**x**_1_, …, **x**_*M*_} and **Y** = {**y**_1_, …, **y**_*M*_}, living in the same Euclidean space ℝ^*N*^, equipped with the standard Euclidean metric:

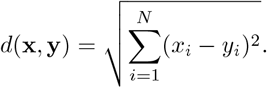

We construct a cost matrix **C** ∈ ℝ^*M ×M*^, where each entry *C*_*ij*_ = *d*(**x**_*i*_, **y**_*j*_)^2^ represents the squared cost of transporting unit mass from **x**_*i*_ ∈ **X** to **y**_*j*_ ∈ **Y**. Let *µ* and *ν* be uniform discrete probability measures over **X** and **Y**, respectively, defined as

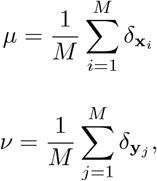

where *δ*_**x**_ denotes the Dirac measure centered at **x**.

The Wasserstein distance of order 2 (also called the Earth Mover’s Distance) is defined as the minimum total cost of transporting *µ* to *ν*, subject to mass conservation:

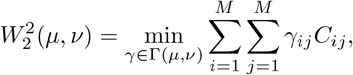

where Γ(*µ, ν*) is the set of joint probability distributions (transport plans) with marginals *µ* and *ν*. In this setting, *γ* represents the optimal transport plan and *γ*_*ij*_*C*_*ij*_ is the amount of probability mass transported from **x**_*i*_ to **y**_*j*_.

The Wasserstein distance provides a natural and geometrically grounded way to compare distributions over vector spaces. However, computing it exactly for empirical datasets can be computationally expensive and numerically unstable, particularly as the number of samples grows. These drawbacks are effectively solved by the concept of Sinkhorn divergence, a regularized version of the Wasserstein distance that preserves its core geometric meaning.

Given two discrete distributions *µ* and *ν* supported on point clouds **X** and **Y**, and a cost matrix **C** as defined above, the entropically regularized transport cost is given by:

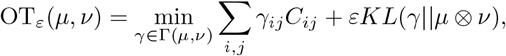

where 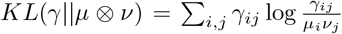 is the Kullback-Leibler penalty that favours factorized optimal plans *γ*, and *ε >* 0 is a regularization parameter controlling the trade-off between accuracy and smoothness. However, this regularized transport cost is not a proper metric: it does not satisfy symmetry or the triangle inequality, and it may not vanish when *µ* = *ν*. The Sinkhorn divergence resolves this by symmetrizing and correcting the regularized cost:

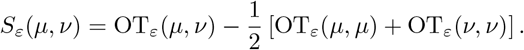

This expression ensures that *S*_*ε*_(*µ, ν*) = 0 if and only if *µ* = *ν*, and retains the geometry of the Wasserstein space while benefiting from better numerical stability and computational efficiency. The Sinkhorn divergence can be efficiently computed using iterative matrix scaling algorithms (e.g., Sinkhorn–Knopp) and is particularly well-suited to comparing empirical distributions like the PCA projections of natural and generated sequences [26].

In our analysis, we use the Sinkhorn divergence with squared Euclidean cost and uniform weights over the empirical samples, as implemented in the Julia OptimalTransport.jl library, setting the regularization parameter *ϵ* = 0.05.

### F Principal components higher than the second

Traditional autoregressive and energy-based models such as ArDCA and bmDCA are trained without any explicit access to global, low-dimensional features of the sequence distribution. While they often reproduce the marginal statistics and low-rank structure of the MSA, they cannot selectively capture higher-order variation, especially along principal components beyond the first few. In practice, this means that although these models can approximate the dominant modes of variation, typically PC1 and PC2, they fail to reproduce finer, structured diversity in the MSA, which is often functionally or structurally meaningful.

In contrast, FeatureDCA incorporates a tunable number of principal components as conditioning inputs, allowing the model to explicitly learn and generate along increasingly complex directions of evolutionary variation. This flexibility enables FeatureDCA to match the natural sequence distribution not just along the first few PCs, but also across mid- and higher-rank components, depending on the dimensionality *d* used during training.

Figure S1 illustrates this effect for family PF13354. It compares the projections of natural and generated sequences across the first ten principal components (shown pairwise as PC1 vs PC2, PC3 vs PC4, …, PC9 vs PC10). Sequences generated by ArDCA and bmDCA align well with the natural data only in the first projection, while diverging noticeably in higher PC pairs. In contrast, FeatureDCA, when trained with increasing numbers of PCs (*d* = 2, 4, 8, 16), shows progressively improved agreement across the full range of components, demonstrating its ability to capture structured variability at different scales.

### A GIn-silico Deep Mutational Scanning

To evaluate the ability of FeatureDCA to predict mutational effects, we performed an in-silico deep mutational scanning (DMS) experiment. The predictive score for a mutation at position *i*, from amino acid *a* to amino acid *b*, is computed as the difference in log-likelihood between the mutant and the wild-type sequence:

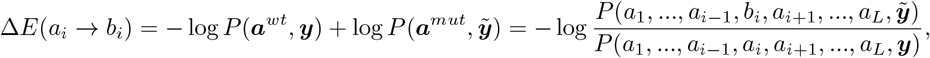

where ***a***^*wt*^ is the wild-type sequence and ***a***^*mut*^ is the same sequence with residue *a* at position *i* replaced by residue *b*, while ***y*** and 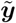 are the PC projections of the wild-type and mutant sequence, respectively. The probabilities *P* (***a, y***) are obtained directly from the model’s autoregressive probability decomposition.

The experimental DMS measurements used for comparison were taken from the Excel file available from https://static-content.springer.com/esm/art%3A10.1038%2Fs41592-018-0138-4/MediaObjects/41592_2018_138_MOESM4_ESM.xlsx[7]. We used the data corresponding to the blat ecoli dataset, which refers to the TEM-1 beta-lactamase experiment by Ostermeier and collaborators [36]. To ensure consistency between model predictions and experimental values, we preprocessed the Ostermeier dataset as follows. First, we alphabetically ordered the amino acid substitutions at each site (A, C, D, E, …, Y) for reproducibility. For any missing single-point mutation, we explicitly added an entry with a placeholder mutational score of inf, representing absence of data. We then constructed a multiple sequence alignment (MSA) for PF13354 with a fixed length of 214 positions, including 7 alignment gaps, and ensured that the wild-type sequence used in the original Ostermeier experiment was included in the alignment. Data relative to the study of the DMS of Beta-Lactamase is available at https://github.com/francescocaredda/FeatureDCAData This processed MSA was used to train FeatureDCA and to compute the in-silico mutational scores. When comparing to the experimental DMS data, we excluded mutations involving missing data (i.e., those with an inf label) as well as trivial identity mutations (e.g., *a* → *a*), which would artificially inflate correlation but carry no functional meaning.

**Figure S1:**
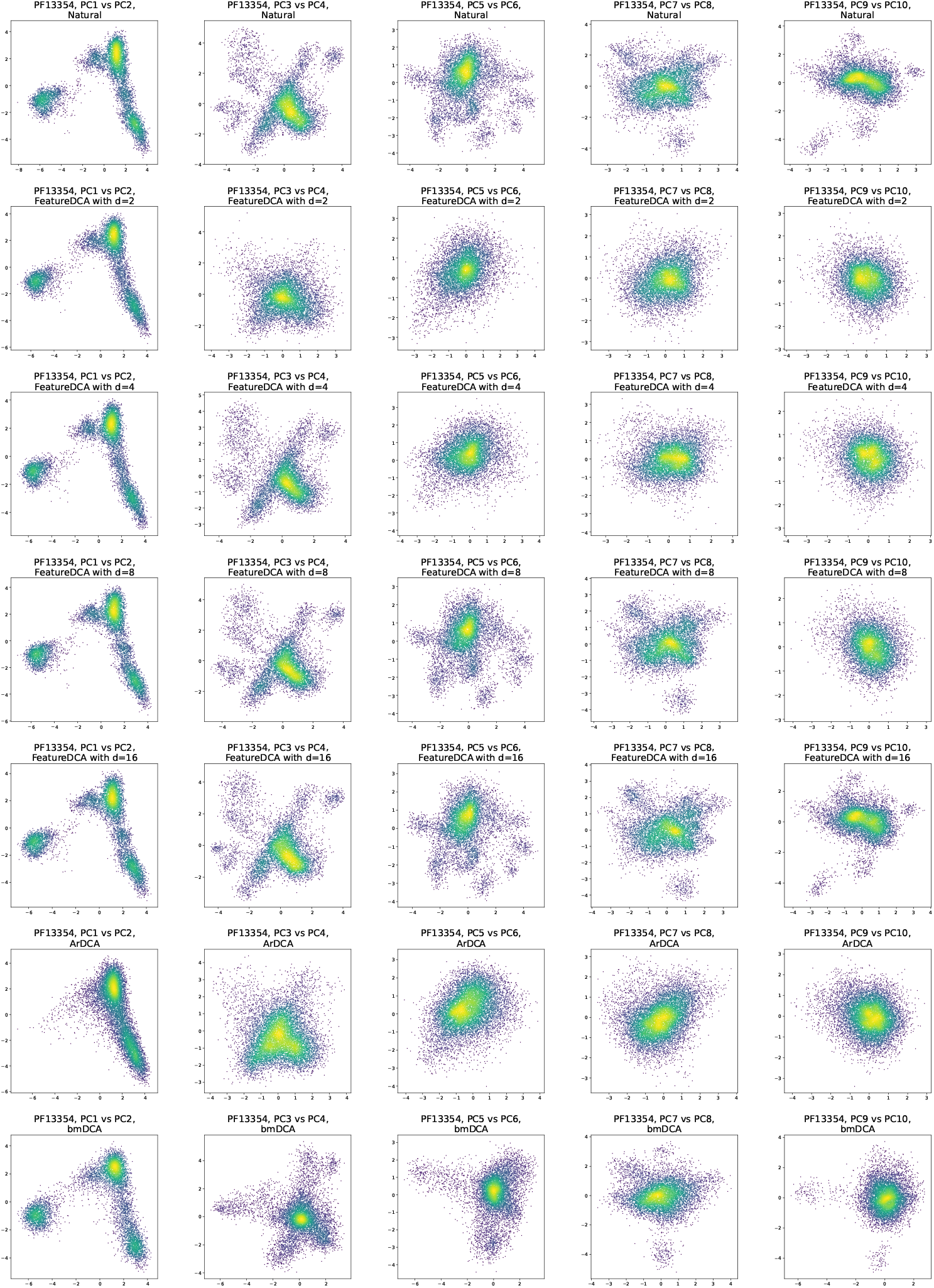
Principal component projections of natural and generated sequences for PF13354. Each column corresponds to a projection onto a pair of principal components: PC1 vs PC2, PC3 vs PC4, …, up to PC9 vs PC10. The first row shows natural MSA sequences. Subsequent rows display sequences generated by different models: FeatureDCA trained with *d* = 2, 4, 8, 16 principal components, ArDCA, and bmDCA. The figure illustrates that FeatureDCA trained with *d* PCs is able to faithfully reproduce the natural sequence distribution within the corresponding first d principal components, with progressively improved agreement as *d* increases.

Figure S2 shows the Spearman rank correlation between the predicted and experimental mutational effects as a function of the number of principal components *d* used to condition FeatureDCA. For small values of *d*, FeatureDCA performs comparably to ArDCA [5]. As *d* increases, a nonmonotonic behavior is observed: the correlation initially drops, then recovers for higher values of *d*, indicating a balance between representational power and overfitting.

**Figure S2:**
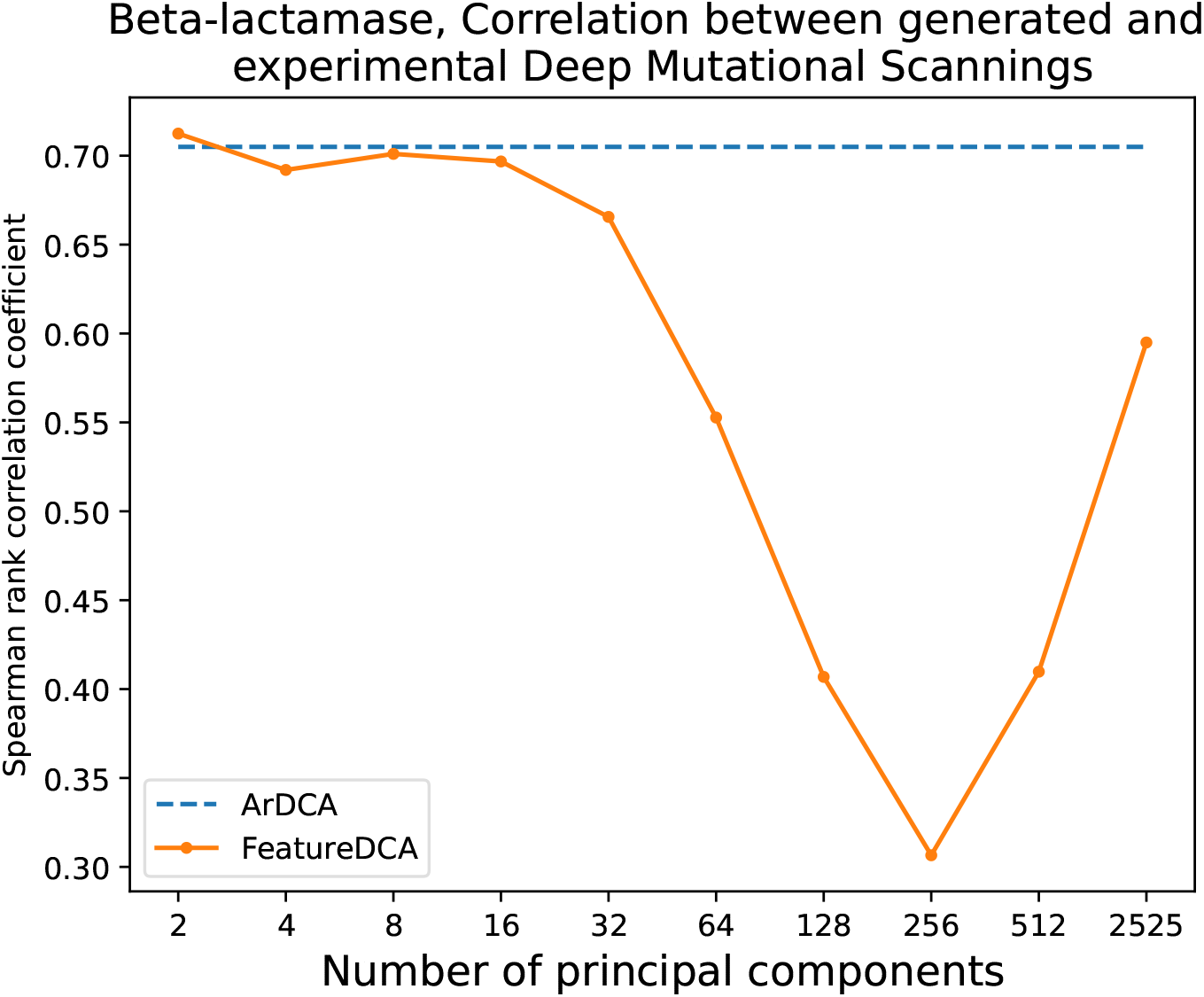
Predictive performance of FeatureDCA and ArDCA on experimental deep mutational scanning data for Beta-lactamase (PF13354). The Spearman rank correlation coefficient is shown between predicted and experimental mutational effects as a function of the number of principal components used during training and generation. ArDCA corresponds to the unconditioned autoregressive baseline. FeatureDCA performs comparably to ArDCA for small values of the number of principal components *d*, but displays a non-monotonic behavior: the correlation initially decreases as more components are introduced, then increases again at larger *d*, suggesting a complex trade-off between dimensionality, signal strength, and generalization.

### H Supplementary Figures

The following figures complement the main text analysis on conditioned generativity. While the main text focuses on family PF13354, the figures below report analogous analyses for four additional protein families: PF00014, PF00072, PF00076, and PF00595. These include principal component (PC) projections, pairwise correlation statistics, effective sequence diversity, and structural fidelity measures for sequences generated by FeatureDCA under different conditioning schemes.

- **Figure S3**: Pearson correlation between connected pairwise correlations (natural vs. generated) and Effective depth (sequence variability) of generated MSAs as a function of the number of principal components used during training and generation. These results correspond to those presented for PF13354 in Figure 2C-D) of the main text
- **Figure S4**: Wasserstein distances between the PCA distributions of natural and generated sequences for different models, using Sinkhorn divergence as described in Supplementary Section D. These results parallel the PCA-matching trends shown in Figure 2B) of the main text.
- **Figure S5**: Projection of natural and generated sequences in the PCA space. This figure extends the PC projection plots shown for PF13354 in 2A) of the main text.
- **Figure S6**: Hamming distance and PC distance between generated and natural sequences for the four families, complementing the analysis of sequence diversity shown in Figure 3 of the main text for PF13354.
- **Figure S7**: Structural distance between generated and experimental wildtype structures for the four families, complementing the analysis of structural accuracy shown in Figure 4 of the main text for PF13354.

**Figure S3:**
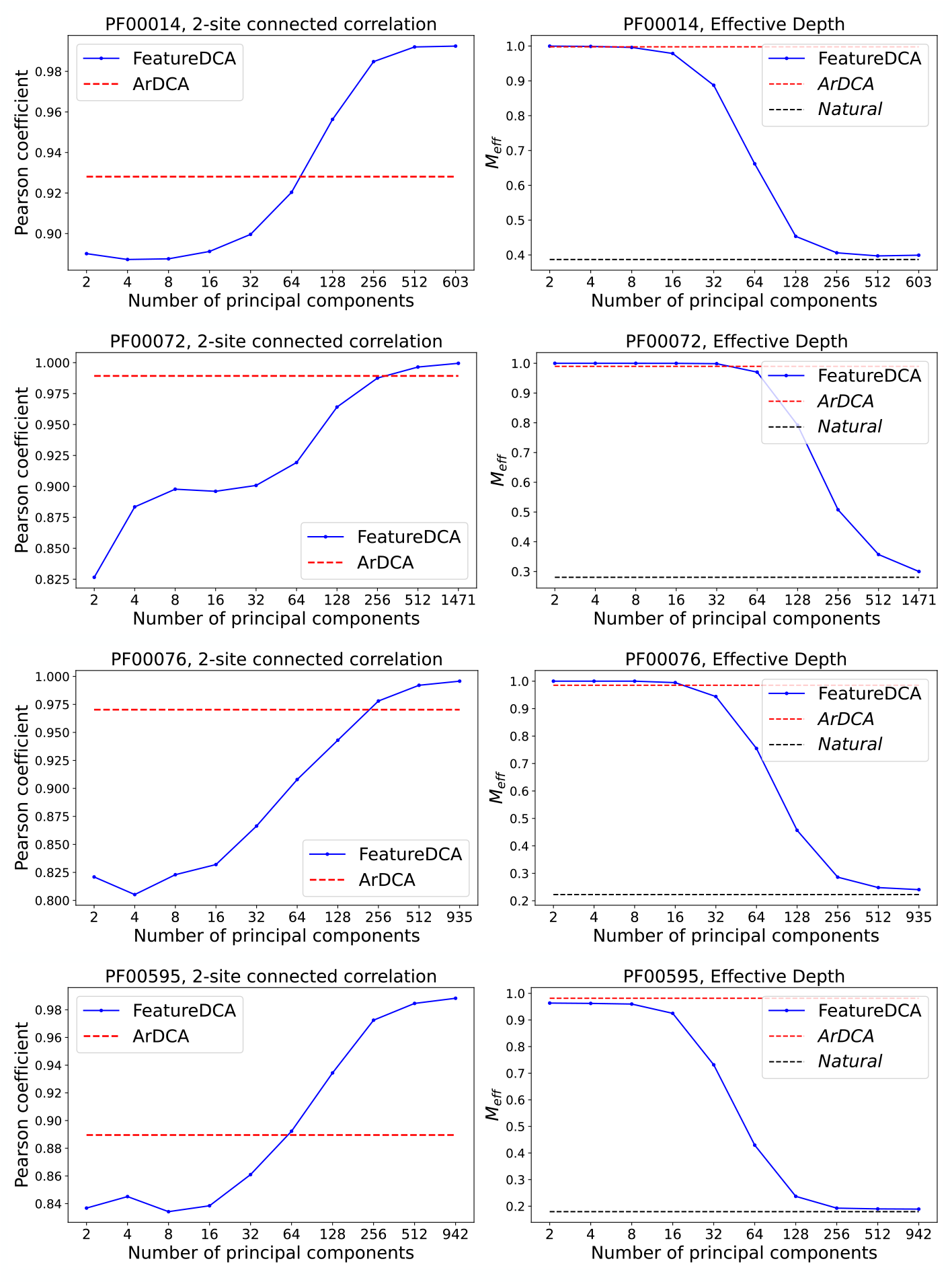
Comparison of generative and statistical properties across protein families as a function of PCA feature dimensionality. Each row corresponds to a different protein family (PF00014, PF00072, PF00076, PF00595). **Left column**: Pearson correlation between the pairwise connected correlations computed on natural sequences generated by FeatureDCA, plotted as a function of the number of principal components used during training and generation (blue line). The red dotted line indicates the baseline correlation obtained from ArDCA. **Right column**: Effective depth (i.e., sequence diversity) of generated datasets as a function of the number of principal components (blue line: FeatureDCA). The red dotted line shows the ArDCA baseline, and the black dotted line marks the effective depth of the natural MSA.

**Figure S4:**
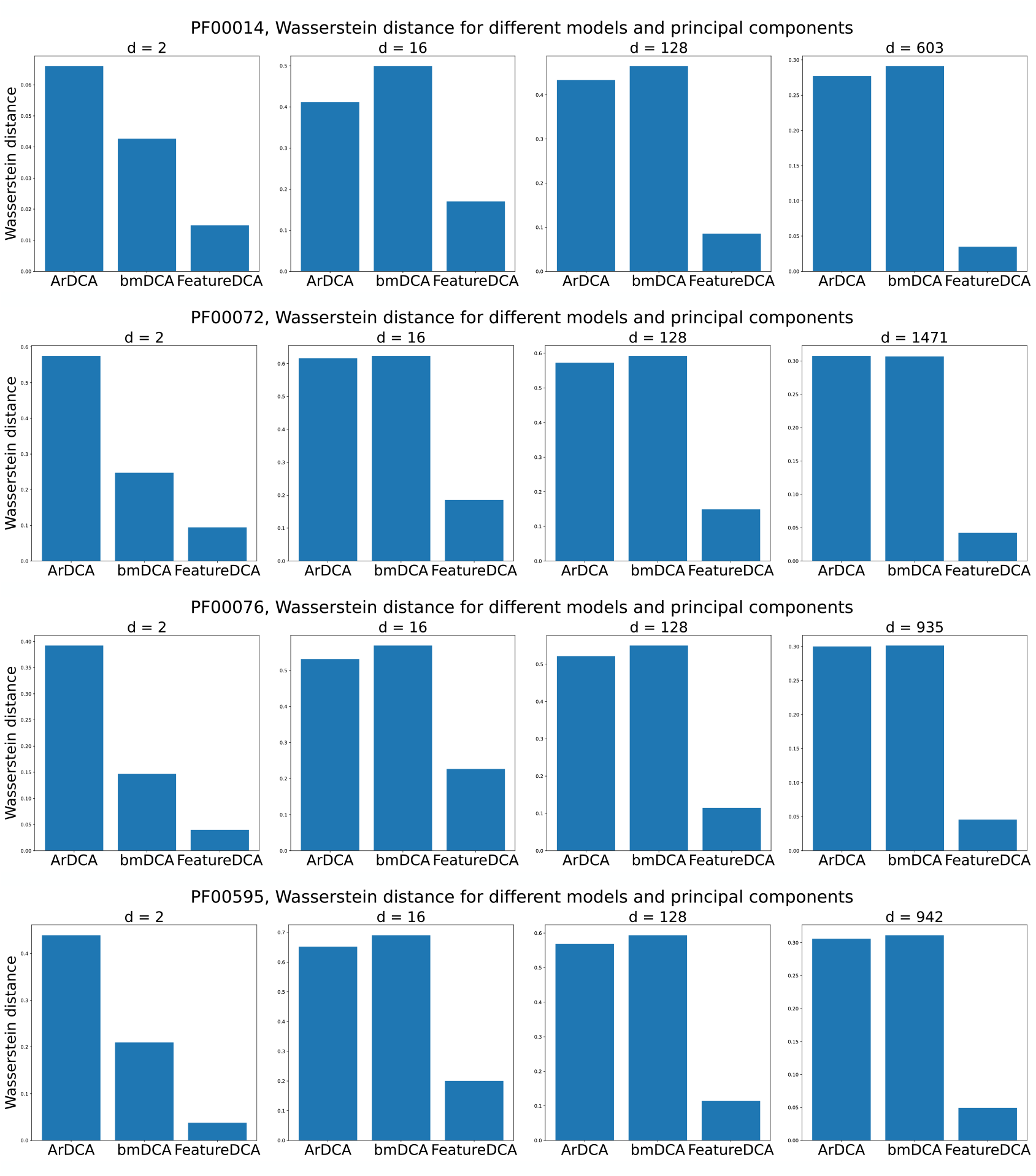
Histogram showing the Wasserstein distance between the *d*-dimensional PCA distributions of the natural and generated MSA for different models. Each row corresponds to a different protein family (PF00014, PF00072, PF00076, PF00595). FeatureDCA was trained with the corresponding number of principal components used to compute the Wasserstein d-dimensional distance.

**Figure S5:**
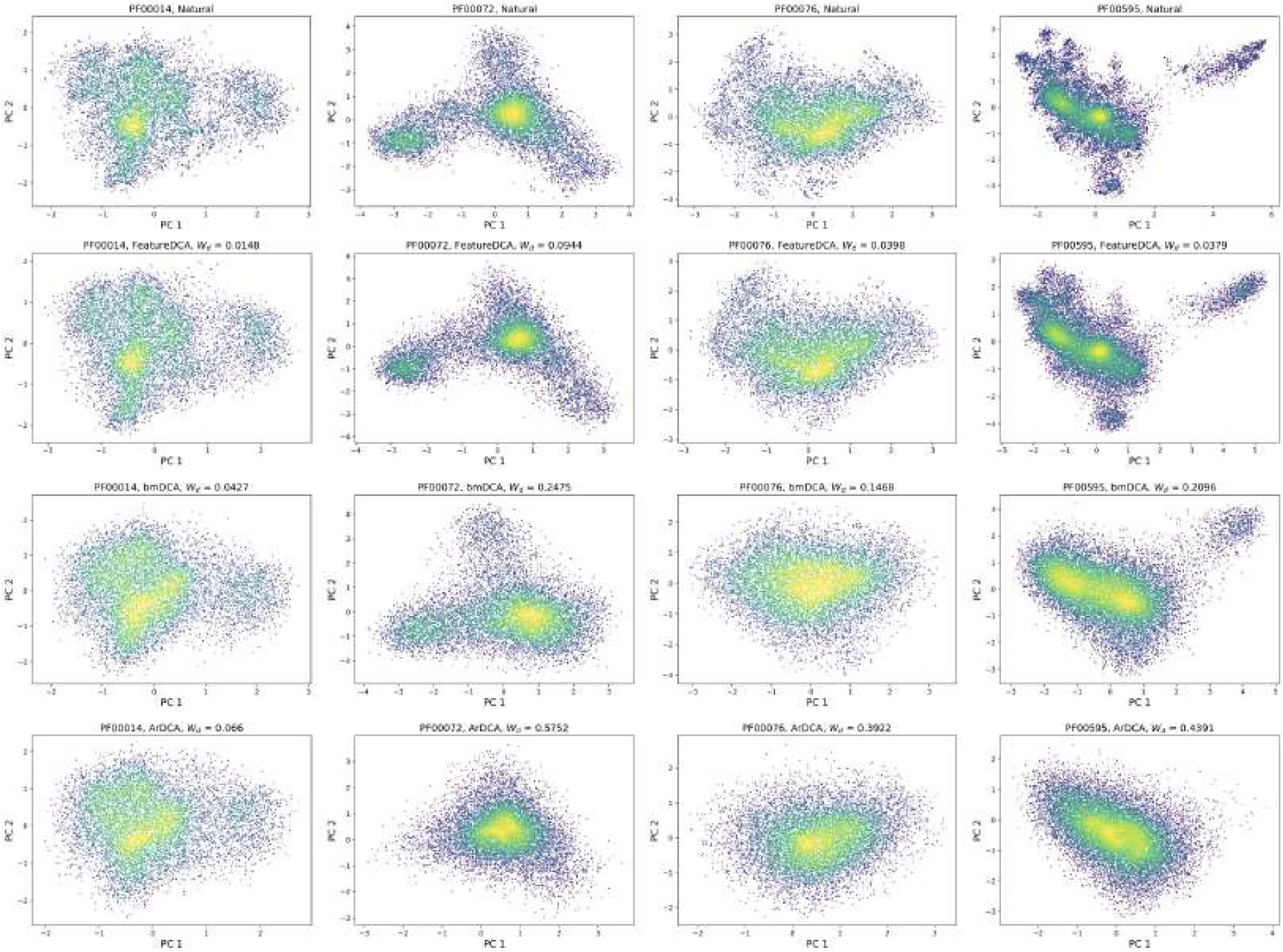
Projection onto the first two principal components of natural and generated MSAs for different protein families. Each column corresponds to a different protein family (PF00014, PF00072, PF00076, PF00595). The first row shows the PCA projection of natural sequences, while the subsequent rows show the projections of sequences generated by FeatureDCA, bmDCA, and ArDCA, respectively. For each model and family, the Wasserstein distance *W*_*d*_ quantifies the discrepancy between the natural and generated distributions along the first two principal components.

**Figure S6:**
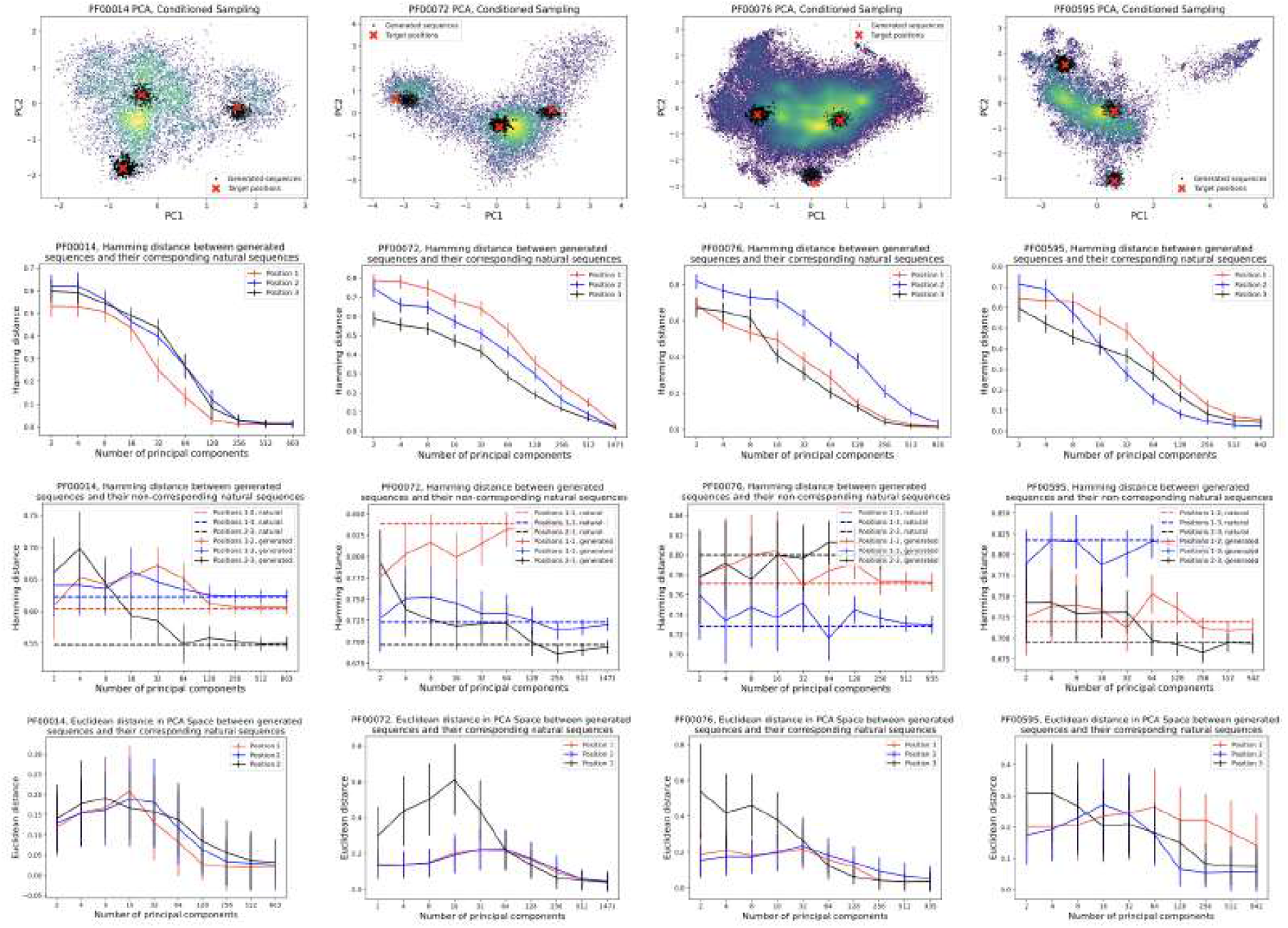
Statistical analysis of the generativity conditioned on three different positions on the PC space as a function of the number of principal components learned during training. Each column corresponds to a different protein family (PF00014, PF00072, PF00076, PF00595). **First row**: red crosses represent the three positions chosen on different islands of the PC projection to study the different conditioned sampling. The clouds of black dots represent the generated sequences around the target positions. **Second row**: Hamming distance between the target positions and the generated sequences conditioned on those positions. **Third row**: Hamming distance between the generated sequences and the non-corresponding target positions. **Forth row**: Euclidean distance in the first two PC plane between the target positions and the generated sequences conditioned on those positions.

**Figure S7:**
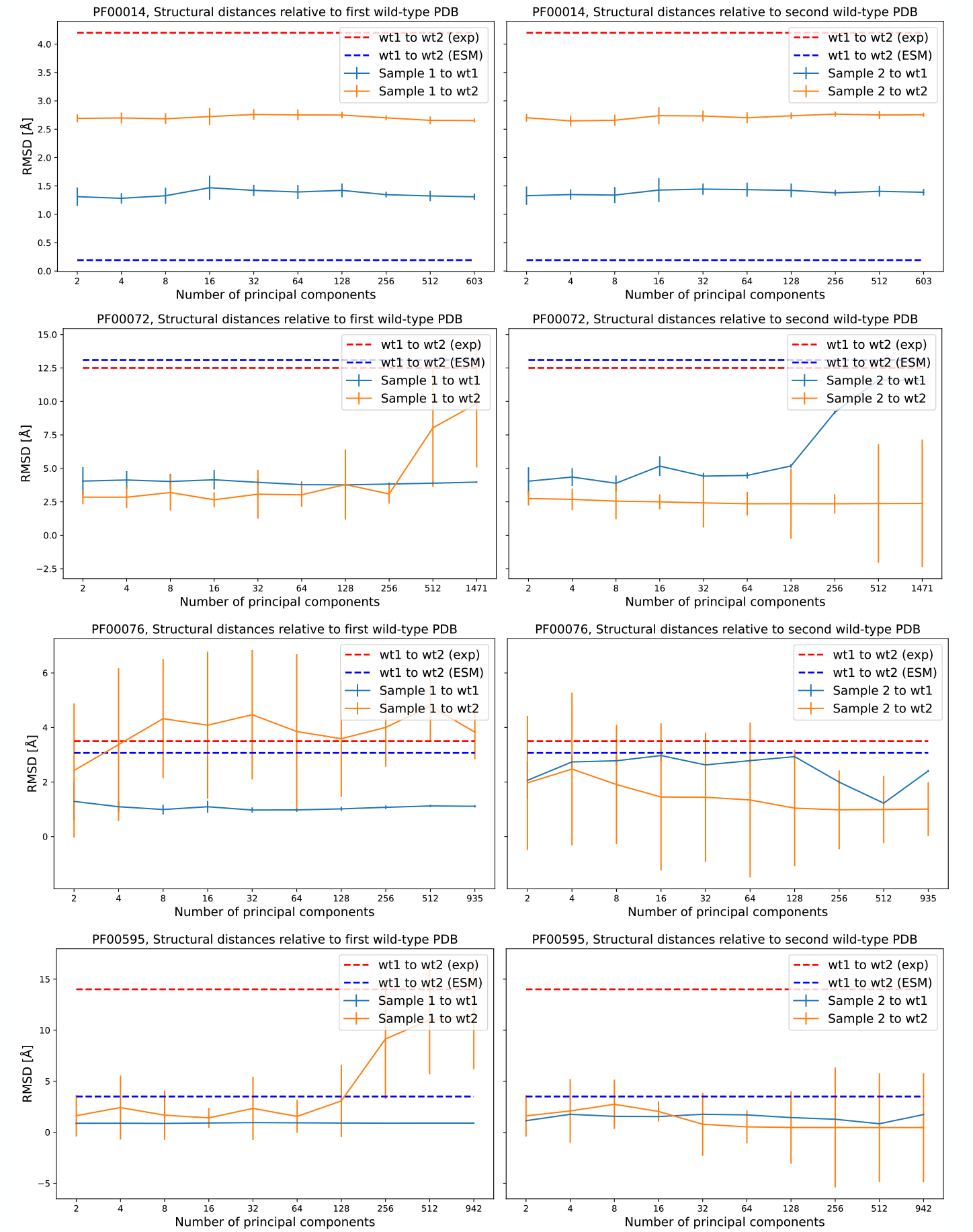
Structural similarity of generated sequences to wild-type structures for different protein families. Each row corresponds to a different protein family (PF00014, PF00072, PF00076, PF00595). Each panel shows the RMSD (Å) between the ESM predicted structures of generated sequences and their respective wild-type references, plotted against the number of principal components used for learning and sequence generation. **Left**: RMSD relative to the first wild-type (wt1) structure; **Right**: RMSD relative to the second wild-type (wt2) structure. Dashed lines indicate the RMSD between the two wild types using experimental structures (red) and ESM predictions (blue). Solid lines represent average RMSDs of generated samples to wt1 (blue) and to wt2 (orange), with error bars showing standard deviation.

## Notes

https://github.com/francescocaredda/FeatureDCAData

https://github.com/francescocaredda/FeatureDCA.jl

### Competing Interest Statement

The authors have declared no competing interest.

